# Iron addicted colorectal cancers exploit Heme-Complex II axis to resist oxidative cell death

**DOI:** 10.1101/2025.11.05.686813

**Authors:** Chesta Jain, Muqit Essani, Roshan Kumar, Nupur K. Das, Rashi Singhal, Nicholas J. Rossiter, Brandon Chen, Wesley Huang, Zheng Hong Lee, Sumeet Solanki, Yuezhong Zhang, Peter Sajjakulnukit, Li Zhang, Prarthana J. Dalal, David A Hanna, Shannon McCollum, Elena M. Stoffel, Joel K. Greenson, L James Maher, Costas A. Lyssiotis, Ruma Banerjee, Yatrik M. Shah

## Abstract

Colorectal cancer (CRC) cells are addicted to iron, which fuels nucleotide synthesis, mitochondrial respiration, and rapid proliferation. Yet paradoxically, high intracellular iron is cytotoxic to most other cells, raising the question of how CRC cells tolerate and exploit iron-rich environments. One pathway thought to mediate iron toxicity is ferroptosis, an iron-dependent form of cell death. However, most ferroptosis regulators were identified through synthetic chemical screens or small molecule activators, and it remains unclear whether these canonical pathways explain how iron itself triggers cell death, particularly *in vivo*. Here, using multi-omics profiling, CRISPR screening, and in vivo models, we uncover a heme–succinate dehydrogenase (SDH)–Coenzyme Q (CoQ) axis that enables CRC cells to buffer iron-induced oxidative stress. Heme-dependent SDH reduces CoQ, which redistributes to mitochondrial and plasma membranes to detoxify lipid ROS as a radical trapping antioxidant. This pathway functions alongside, and in some contexts independently of, canonical ferroptosis regulators. These findings reveal that CRCs co-opt metabolic cofactors not only for growth but also for survival under physiologically toxic iron levels, uncovering new vulnerabilities for therapy.

## INTRODUCTION

Colorectal cancer (CRC) is the third most diagnosed cancer globally and the third leading cause of cancer related mortality^1^. CRC cells are remarkably resilient, displaying profound metabolic adaptations that enable survival and growth in nutrients and oxygen deficient microenvironments^2^. This metabolic rewiring is essential for optimal nutrient utilization and prioritizing cellular energetics for proliferation^2^. While such adaptations support tumor growth, they also create dependencies on specific metabolites and micronutrients. Although much attention has been focused on the role of altered metabolism in tumorigenesis, the contribution of micronutrients to cancer progression remains poorly understood.

Iron is a critical micronutrient required by nearly all cells. Both systemic and cellular iron homeostasis are tightly regulated through transcriptional and translational control of iron uptake, utilization, and storage^3^. Dysregulation of these mechanisms can lead to iron deficiency or iron overload, manifesting clinically as anemia or hemochromatosis, respectively^4^. At the cellular level, labile iron supports proliferation, ATP production, and mitochondrial respiration through its redox activity. The role of iron and its regulated pathways in cancer initiation, progression, and metastasis is well established and continues to be a highly active area of investigation^5^.

In CRC, epidemiological studies have linked elevated systemic iron, with increased CRC incidence^6–8^. CRC cells accumulate significantly higher levels of iron compared to adjacent normal tissue, largely by enhancing iron uptake and limiting export^9–12^. We and others have shown that restricting iron availability reduces proliferation, lowers tumor burden, and improves survival in mouse models^9,12^. However, therapeutic iron depletion remains clinically challenging. CRC cells can access iron from both dietary sources and systemic circulation, and lowering intracellular iron primarily slows proliferation rather than eliminating transformed cells^9,13^. Excess intracellular iron is toxic, driving a form of lipid peroxidation–dependent, non-apoptotic cell death known as ferroptosis^14,15^. This creates a striking paradox; CRC cells sequester and accumulate iron and resist its toxicity. Previous work has pointed to elevated antioxidant defenses in CRC models and patient tissues, but the specific metabolic mechanisms that allow cancer cells to tolerate such iron overload remain unknown^16^. This represents a fundamental gap in our understanding of CRC biology. How do CRC cells withstand and even thrive under conditions that should be lethal due to iron-induced oxidative stress? Addressing this question is critical, as uncovering these adaptive mechanisms could expose new therapeutic vulnerabilities.

We show that the canonical anti-ferroptosis response mediated by GPX4 and SLC7A11 is dispensable for CRC growth and progression in both sporadic and colitis-associated cancer models. A metabolism focused CRISPR screen revealed a protective role of cellular heme against iron toxicity. As a critical cofactor for mitochondrial electron transport chain (ETC) enzymes, heme depletion caused profound mitochondrial defects, including loss of complex II (CII) activity. Through multi-omics profiling of *in vitro* and *in vivo* CRC models, we identify a previously unrecognized role of mitochondrial CII in buffering iron-induced cell death by regulating coenzyme Q (CoQ) biosynthesis. Strikingly, pharmacological, or genetic inhibition of CII sensitized CRC cells to iron-induced toxicity *in vitro* and *in vivo*. Collectively, these findings uncover a non-canonical antioxidant function of CII in protecting CRC cells from iron-induced toxicity. By defining how CRC cells adapt to survive under iron overload, our study addresses a major gap in the field and highlights metabolic strategies for development of new therapies.

## RESULTS

### GPX4 is dispensable for CRC initiation and progression

A defining paradox in CRC biology is its ability to accumulate and tolerate exceptionally high levels of intracellular iron^9^. Iron is indispensable for DNA synthesis, mitochondrial metabolism, and rapid proliferation^13,17^. Yet in most cells, sustained iron loading is toxic, as it accelerates reactive oxygen species (ROS) production and triggers ferroptosis, an iron-dependent cell death program driven by lipid peroxidation^14,15^. The prevailing paradigm is that cancer cells buffer this stress primarily through the GPX4–SLC7A11 antioxidant axis, which detoxifies lipid peroxides at the expense of glutathione^18,19^. However, whether this canonical ferroptosis defense is required for CRC initiation or progression has not been directly addressed.

We first assessed the expression of *SLC7A11* and *GPX4* in tumor tissues and matched normal mucosa from individual CRC patients, and in enteroids generated from these same tumors and matched normal tissues. In all cases, RNA transcripts for both were significantly elevated in the tumor-derived material compared to matched normal controls (Fig. 1A–B), indicating that upregulation of these pathways is intrinsic to the tumor epithelium and maintained *ex vivo*. These findings were consistent with TCGA datasets, where *GPX4* and *SLC7A11* transcripts are elevated in CRC tumors relative to normal colon tissue (Fig. S1A). Despite this consistent upregulation, Kaplan–Meier survival analysis revealed no correlation between high *GPX4/SLC7A11* expression and overall patient survival (Fig. 1C), suggesting that these pathways may not be essential for CRC fitness.

**Figure 1:**
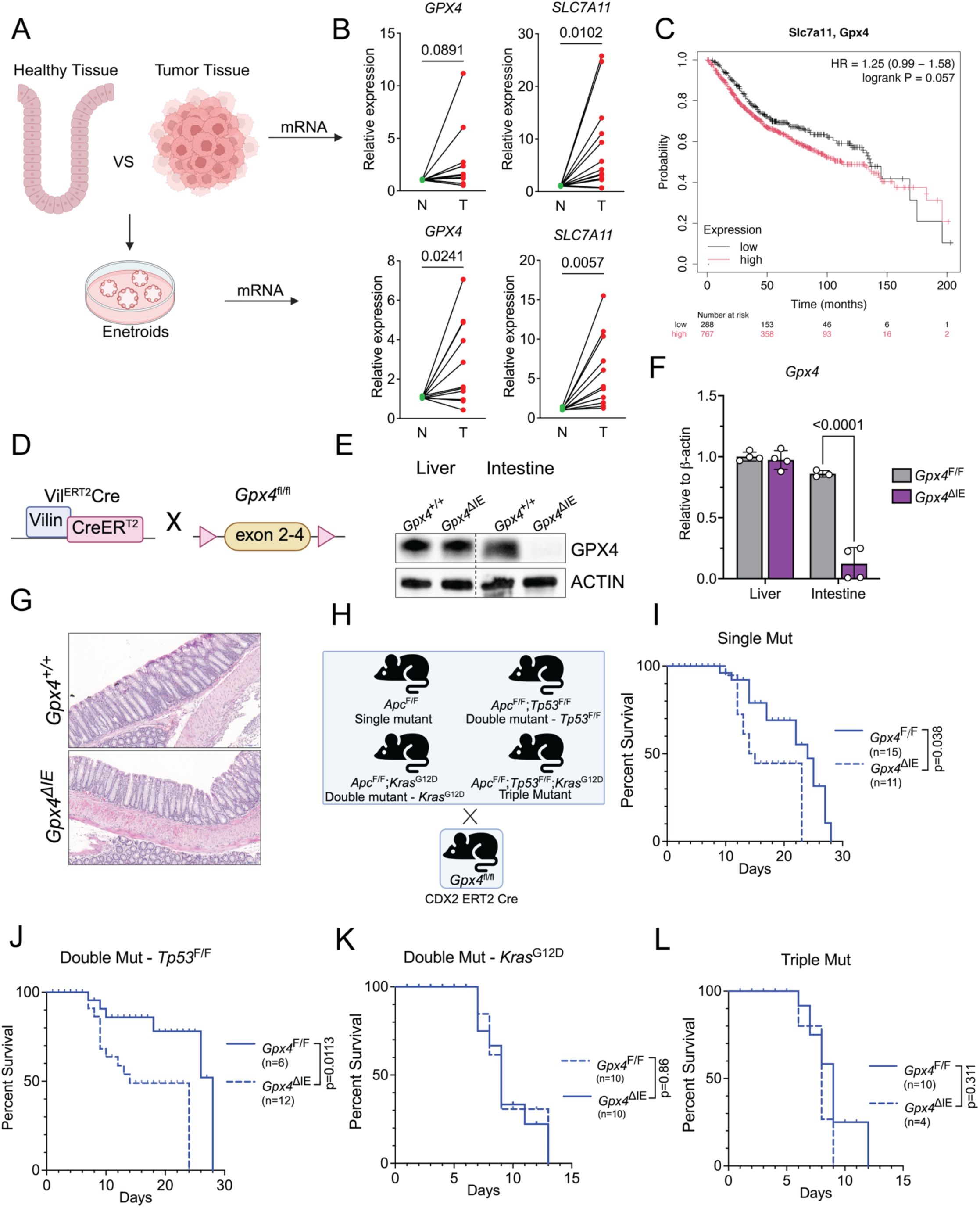
GPX4 is Dispensable for CRC Initiation and Progression. A) Schematic showing enteroid isolation from normal and tumor tissues from patient biopsies. B) qPCR showing mRNA expression of *GPX4* and *SLC7A11* from patient derived tumor and normal adjacent tissues and enteroids. C) Kaplan-Meier survival curves of colon adenocarcinoma cancer patients stratified by the *GPX4* and *SLC7A11* gene expression with data from TCGA cancer database. D) Schematic showing breeding strategy to generate tamoxifen inducible intestine specific GPX4 deficient mice. E) Western blot showing hepatic and intestinal GPX4 protein levels in GPX4^+/+^ and GPX4^ΔIE^ mice 5 days post tamoxifen induction via 100 mg/kg body weight I.P. injection. F) qPCR of colonic GPX4 transcript levels in GPX4^+/+^ and GPX4^ΔIE^ mice from (E). G) H&E of GPX4^+/+^ and GPX4^ΔIE^ mice from (E) showing normal colonic histology. H) Schematic showing mouse model breeding strategy to generate Single/Double-*Tp53*/Double-*Kras*^G12D^/Triple mutant mice with or without functional GPX4. I) Survival plot of GPX4 deficient Single J) Double-*Tp53*, K) Double-*Kras*^G12D^ L) Triple mutant mice post tamoxifen induction via 100 mg/kg body weight I.P. injection. All data are mean ± SEM. t tests for (B), (I), (J), (K), (L) or One-way ANOVA with Tukey’s multiple comparisons test (F). ∗p < 0.05, ∗∗p < 0.01, ∗∗∗p < 0.001.

To test this functionally, we generated intestine-specific *Gpx4* knockout mice (Fig. 1D–F). Intestine specific *Gpx4* deletion does not manifest as any significant histological or physiological changes and does not alter overall GI health or overall survival (Fig. 1G)^20^. In the colitis-associated cancer (CAC) model*, Gpx4* loss did not significantly alter tumor number, size, or overall burden (Fig. S1B–E). While these findings suggest that CRC cells rely on metabolic adaptations beyond the canonical SLC7A11-GPX4 dependent anti-ferroptotic response, colitis associated cancer is rare in human population and majority of the sporadic CRC cases are *APC*, *P53* and *KRAS* mutation driven.

To examine the role of GPX4 in sporadic CRC, we crossed the *Gpx4*^F/F^ mice with a series of genetically engineered colon cancer models: Single mutant, (*Apc*^F/F^), Double mutant (*Apc*^F/F^; *Tp53*^F/F^) or Triple mutant (*Apc^F^*^/F^; *Tp53*^F/F^; *Kras*^G12D^)^21^. All mutations were specifically targeted to the colon epithelium using CDX2-CreERT2 (Fig. 1H). If GPX4 were essential to buffer tumoral iron and resist iron-induced toxicity, its deletion would be expected to slow tumor progression and improve survival. Surprisingly*, Gpx4* deficiency did not improve survival in any of these models (Fig. 1I-L). There were no histological differences observed in the colon between the *Gpx4*^F/F^ and *Gpx4*^ΔIE^ (Fig. S1F). Notably, in mice harboring *Apc* or *Apc/Tp53* mutations, *Gpx4* deficiency slightly shortened lifespan, which may reflect altered inflammatory responses in the absence of GPX4^22^. This indicates that CRC does not depend on the canonical SLC7A11-GPX4 ferroptosis defense to tolerate iron stress. Instead, CRC must exploit alternative metabolic adaptations that enable survival under high iron.

### CRC cells resist iron toxicity through metabolic adaptations

To understand if the resistance to high iron levels is a cell autonomous mechanism, we first compared proliferation under millimolar concentrations of ferric ammonium citrate (FAC) in CRC cell lines versus normal intestinal epithelial cells and non-CRC cancer lines. CRC cells maintained robust proliferation, whereas non-CRC cells rapidly succumbed to iron toxicity (Fig. 2A–C). Sensitivity to iron varied across CRC lines, reflecting heterogeneity in intrinsic metabolic adaptations (Fig. S2A). Chronic high-iron exposure reduced CRC growth, sensitized cells to ferroptosis inducers (RSL3 and erastin) and induced cell death, which was partially rescued by ferroptosis inhibitors and antioxidants such as liproxstatin (Lip-1) and N-acetyl cysteine (NAC) (Fig. S2B, Fig. 2D;). By contrast, iron-induced death in non-CRC lines (fibroblasts CD1, NIH3T3, and fibrosarcoma HT1080) was fully reversed by ferroptosis inhibition (Fig. S2C-D).

**Figure 2:**
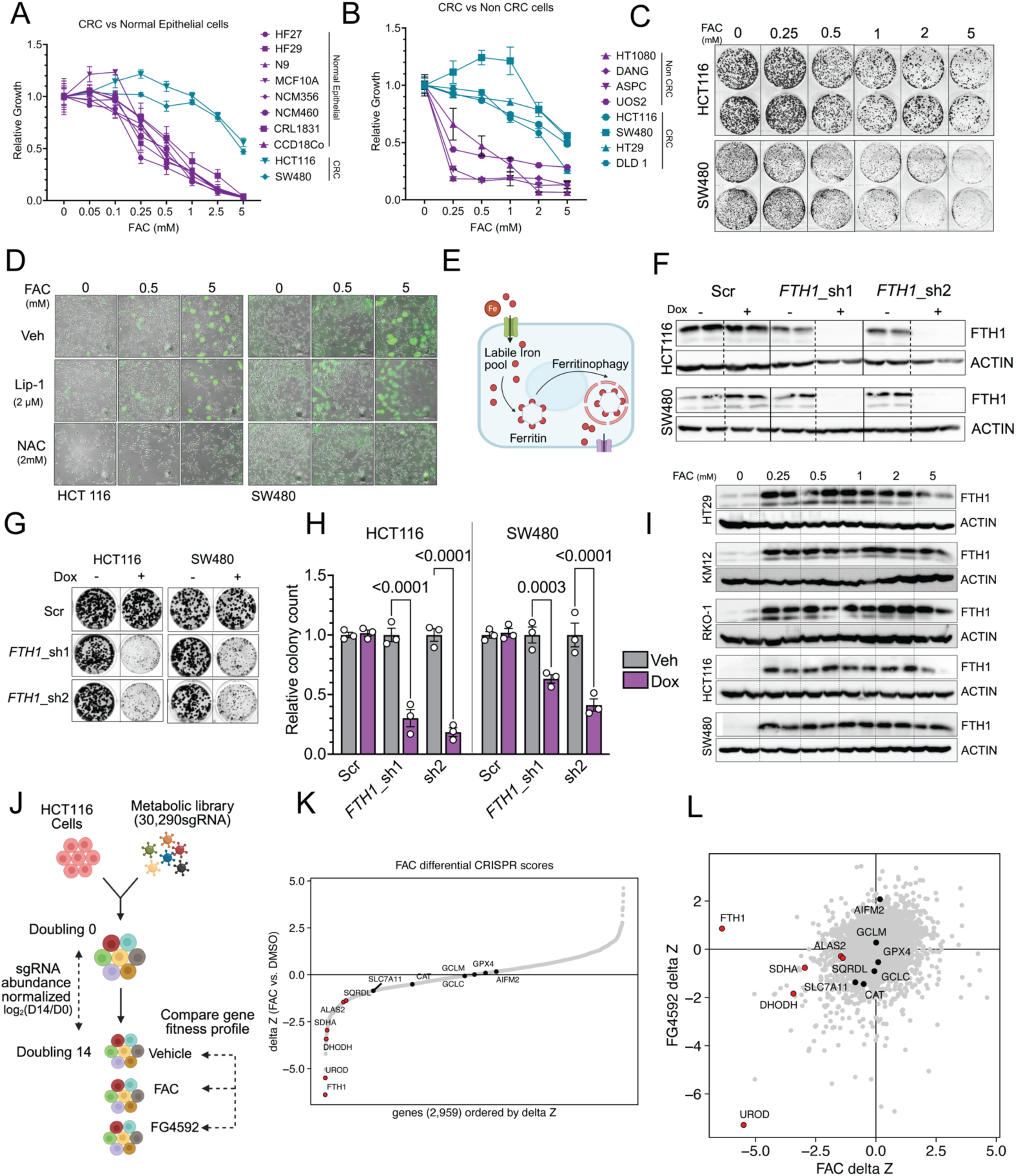
CRC Cells Resist Iron Toxicity Through Metabolic Adaptations. A) Relative growth of CRC or normal epithelial cells 72h post FAC treatment measured using bright field microscopy. B) Relative growth of CRC or non-CRC cancer cells 72h post FAC treatment measured using bright field microscopy. C) Representative crystal violet staining images of HCT116 and SW480 cells 14 days post FAC treatment. D) Representative sytox green staining images from HCT116 and SW480 cells showing dead cells in green 72h post FAC treatment with or without ferroptosis inhibitor Liproxstatin (Lip-1, 2μM) or N-acetyl cysteine (NAC, 5mM). E) Schematic showing role of ferritin in maintaining cellular iron homeostasis. F) Western blot images showing protein expression of FTH1 and beta Actin in HCT116 or SW480 cells stable for empty tet-pLKO-puro (Scr) or shRNA targeting FTH1 (shRNA 1 or shRNA 2) 48h post doxycycline (dox) treatment (250ng/mL). G) Representative crystal violet staining images of cells from (F) 14 days post dox treatment. H) Relative fold change quantified from (G) I) Western blot images showing protein expression of FTH1 and beta actin in CRC cell lines 24h post FAC treatment at different concentrations. J) Schematic showing workflow for the metabolism focused CRISPR library screen under FAC (2.5mM) or FG4592 (100μM) treatment. K) Differential CRISPR score (DMSO-FAC) highlighting significantly depleted genes of interest in doubling 14 populations. L) Scatter plot comparing CRISPR scores under FAC (x axis) or FG4592 (y axis) treatment. Genes of interest highlighted in red. All data are mean ± SEM. One-way ANOVA with Tukey’s multiple comparisons test (H). ∗p < 0.05, ∗∗p < 0.01, ∗∗∗p < 0.001.

This raises the question: what metabolic adaptations allow CRC cells to survive under iron loads that are lethal for normal intestinal epithelial cells or non-CRC cancer cells? Intracellular iron homeostasis depends on coordinated regulation of iron import/export and ferritin-mediated storage^3^. CRC cells have increased expression of iron importers and hepcidin, an iron export repressor, via sustained HIF2α transcriptional activity^9,10,13^. Ferritin acts as both a bioavailable iron reservoir and a buffer against iron overload (Fig. 2E)^23,24^. To test its role in CRC iron tolerance, we generated CRC lines with doxycycline-inducible shRNA targeting ferritin heavy chain (*FTH1*) (Fig. 2F). *FTH1* knockdown was lethal, markedly reducing proliferation, and growth defects were not rescued by lipid ROS scavengers (Fig. 2G–H, S2E). Interestingly, FTH1 protein levels saturated at low FAC concentrations and did not increase further under higher FAC, and overexpression did not improve iron tolerance (Fig. 2I, S2F–G). These data indicate that while FTH1 maintains basal iron homeostasis, it is insufficient to buffer the extreme labile iron pools in CRC.

To uncover additional mechanisms, we performed a metabolism-focused CRISPR-Cas9 screen^25^. Cells were maintained under chronic FAC exposure to mimic iron overload, or under HIF2α activation via FG4592 treatment, which we and others have shown increases intracellular iron levels and susceptibility to iron-induced toxicity (Fig. 2J)^26^. This approach allowed us to probe metabolic pathways critical for CRC survival under physiologically relevant iron stress conditions. The screen identified depletion of several mitochondrial metabolism-related genes, including those involved in purine/pyrimidine biosynthesis, cholesterol biosynthesis, and heme biosynthesis, under both FAC and HIF2α activation (Fig. 2K–L, S2H). *FTH1* was selectively depleted under FAC but not FG4592, consistent with HIF2α-mediated transcriptional regulation of ferritin. These results suggest that CRC survival under high iron depends on mitochondrial and heme-dependent metabolic pathways, beyond canonical ferritin-mediated iron storage.

### Heme biosynthesis buffers iron-induced toxicity

The metabolism-focused CRISPR-Cas9 screen (Fig. 2L) identified uroporphyrinogen decarboxylase (*UROD*), a key enzyme in heme biosynthesis, as a top-scoring gene significantly depleted under both FAC and HIF2α activation (FG4592) treatments. Heme, an iron-containing cofactor, is essential for numerous cellular processes, including mitochondrial respiration, Fe–S cluster biogenesis, and nucleotide synthesis^27^. While dietary heme is a major source, all cells including CRC cells express the full complement of enzymes required for de novo heme synthesis. Increasing intracellular iron elevates cellular heme pools in CRC cells (Fig. 3A–B). To validate the screen, we generated HCT116 and SW480 cells with Tet-inducible knockdown (KD) of *UROD* using two independent shRNAs (shRNA1 and shRNA2), with empty vector (Scr) controls (Fig. S3A). Consistent with the previous findings, depleting intracellular heme genetically by knocking down *UROD* or pharmacologically using succinyl acetone (SA), a competitive inhibitor of delta-aminolevulinate dehydratase (ALAD) sensitizes CRC cells to iron induced toxicity (Fig. S3B, C; Fig. 3C).

**Figure 3:**
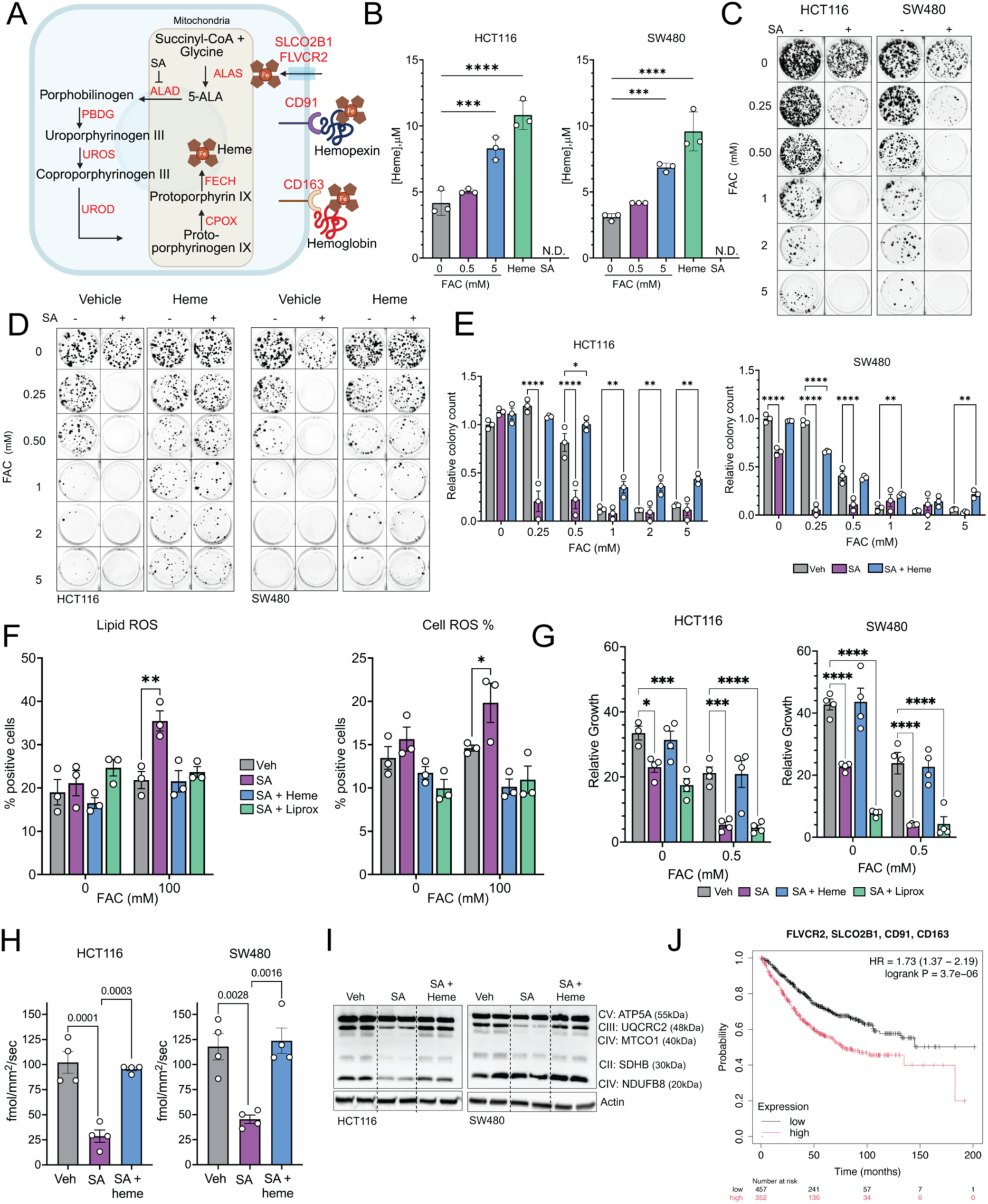
Heme Biosynthesis Buffers Iron-Induced Toxicity. A) Schematic showing cellular heme uptake and biosynthesis pathway in mammalian cells. B) Intracellular heme levels in HCT166 and SW480 cells 72h post FAC (0.5, 5mM), hemin chloride (heme, 10μM) and Succinyl acetone (SA, 500μM) treatment. C) Representative crystal violet staining of cells 14 days post FAC treatment at indicated concentration with or without SA (250μM). D) Representative crystal violet staining of cells 14 days post FAC treatment at indicated concentration with or without SA (500μM) and or heme (10μM). E) Quantification of relative colony count from (D). F) Lipid ROS and Cell ROS measured by C11-BODIPY581/591 and free radical sensor carboxy-H2DCFDA respectively 72h post FAC (100μM) with or without SA (500μM) and heme (10μM). G) Relative growth of HCT116 and SW480 cells 72h post FAC (0.5mM) with or without SA (500μM), heme (10μM) liproxstatin (Lip-1, 2μM). H) Oxygen consumption rate of HCT116 and SW480 cells 72h post FAC treatment (100uM) with or without SA (500μM) or Heme (10μM) measured using the Resipher^TM^ oxygen sensor normalized with cell count measured by brightfield microscopy. I) Western blot images showing expression of subunits of the mitochondrial electron transport chain (CI: NDUFB8, CII: SDHB, CIII: UQCRC2, CIV: MTCO1, CV: ATP5A) in HCT116 and SW480 cells 72h post SA treatment (500uM). J) Kaplan-Meier survival curves of colon adenocarcinoma cancer patients stratified by the FLVCR2, SLCO2B1, CD91, CD163 gene expression with data from TCGA cancer database. All data are mean ± SEM. t tests for (B) or One-way ANOVA with Tukey’s multiple comparisons test for (E), (F), (G), (H). ∗p < 0.05, ∗∗p < 0.01, ∗∗∗p < 0.001.

Heme depletion induced iron sensitivity is completely rescued by supplementing heme back in the culture media (Fig. 3D-E). Knockdown of ferrochelatase (*FECH*), which inserts Fe²⁺ into protoporphyrin IX in the mitochondrial matrix, similarly sensitized CRC cells to iron toxicity, with rescue upon heme supplementation (Fig. S3D–F). These results indicate that heme functions as an essential cofactor supporting antioxidant and metabolic responses, rather than acting as a passive iron store. This response is conserved in other CRC cell lines as well as other cancer lines (Fig. S3G). Moreover, depletion of intracellular heme does not sensitize CRC cells to toxicity associated with other metals or oxidizing agents such as copper, selenium and hydrogen peroxide or the chemotherapeutic agent 5-fluorouracil (5-FU) (Fig. S4A). Interestingly heme supplementation alone conferred resistance to oxidative stress from selenium and H₂O₂, but not to the chemotherapeutic 5-fluorouracil, demonstrating specificity for redox stress (Fig. S4B, C).

Heme depletion increased both general cellular ROS and lipid ROS under FAC treatment, effects that were reversed by either heme supplementation or ferroptosis inhibitor liproxstatin (Lip-1) (Fig. 3F). Interestingly, only heme supplementation rescued SA-induced growth suppression, indicating a non-ferroptotic, heme-dependent oxidative vulnerability (Fig. 3G). This data shows that iron induced toxicity is aggravated by induction of general ROS and lipid ROS; however, quenching these species is not sufficient to reverse iron toxicity, suggesting a non ferroptotic iron-dependent oxidative cell death. Heme depletion also reduced oxygen consumption rates (OCR) and decreased protein levels of ETC complexes (CII, CIII, and CIV) in CRC cells; both parameters were restored by heme supplementation (Fig. 3H, I). These findings are consistent with the known role of heme in mitochondrial respiration and suggest a protective role of heme for buffering iron-induced oxidative stress.

Overall our findings align with the epidemiological data that high expression of genes involved in heme uptake and absorption correlates with reduced overall survival in CRC patients, consistent with the role of heme in promoting tumor cell fitness under high iron conditions (Fig. 3J)^8^. Collectively, these data demonstrate that increase in cellular heme through biosynthesis is a key metabolic adaptation that enables CRC cells to tolerate high intracellular iron, bridging the gap between iron accumulation and oxidative stress resistance.

### Mitochondrial complex II protects CRC cells from iron-induced oxidative cell death

Heme itself can be degraded by hemoxygenase (HO)-1 into the byproducts labile iron, bilirubin and biliverdin^28^. The latter two are hydrophobic molecules with potent antioxidant properties in membranes, where they can protect cells from iron-induced toxicity^29^. However, this process simultaneously releasees free iron, complicating the interpretation of their protective effects. We did not see enhanced iron sensitivity upon alteration of cellular bilirubin and biliverdin levels by inhibition of HO-1 or protection by directly supplementing with products of HO-1 (Fig. S5A-C).

Beyond providing structural and catalytic support to mitochondrial proteins, heme availability has been reported to regulate antioxidant responses through BACH1 (BTB and CNC homology 1) and cap-dependent translation via the heme-regulated kinase(HRI)^30,31^. Free heme can bind to BACH1, a translational repressor of other master antioxidant response transcription factors such as Nuclear factor erythroid 2-related factor 2 (NRF2)^32^. We did not see a significant reduction in the mRNA transcripts of NRF2 targets in SA treated cells (heme deficient) (Fig. S5D, E). Heme deficiency activates HRI kinase activity through autophosphorylation, triggering the ATF4-dependent integrated stress response and reduces translation of mitochondrial proteins including the ETC complexes^31^. We observed no changes in canonical ATF4 target gene expression across FAC, FAC + SA, and FAC + SA + heme treatments (Fig. S5F, G). Notably, mRNA levels of both CHOP and ATF4 were increased in FAC + SA treatment. Since CHOP is broadly induced during cell death, this increase is likely reflective of enhanced cell death rather than a specific heme-regulated signaling effect.

Because ∼20% of total heme-containing proteins function in the ETC and ∼50% serve catalytic roles in diverse metabolic reactions, we next investigated whether heme deficiency disrupts CRC metabolic programs beyond respiration^33^. Our prior work demonstrated that CRC cells rely on iron-dependent nucleotide synthesis, but it was unclear whether this phenotype was strictly iron- or heme-dependent^13^. Metabolomics revealed that SA-induced heme depletion caused a broad metabolic reprogramming, which was only partially corrected by heme supplementation (Fig. 4A). TCA and glycolytic intermediates including lactate, fructose-1,6-bisphosphate (F-1,6BP), glucose-6-phosphate (G6P), and dihydroxyacetone phosphate (DHAP) were markedly reduced in SA-treated cells and restored upon heme repletion, while succinate as accumulated in SA treated cells (Fig. 4B). Under nutrient and oxygen limiting conditions, resembling the tumor microenvironment, amino acids such as glutamine can fuel the TCA cycle to maintain cell metabolism independent of oxygen driven mitochondrial respiration^34,35^. Consistent with this, glutamine, and α-ketoglutarate, as well as glutaminolysis-dependent intermediates including aspartate, orotate, and uridine, accumulated in both SA- and SA+heme–treated cells (Fig. 4B– D).

**Figure 4:**
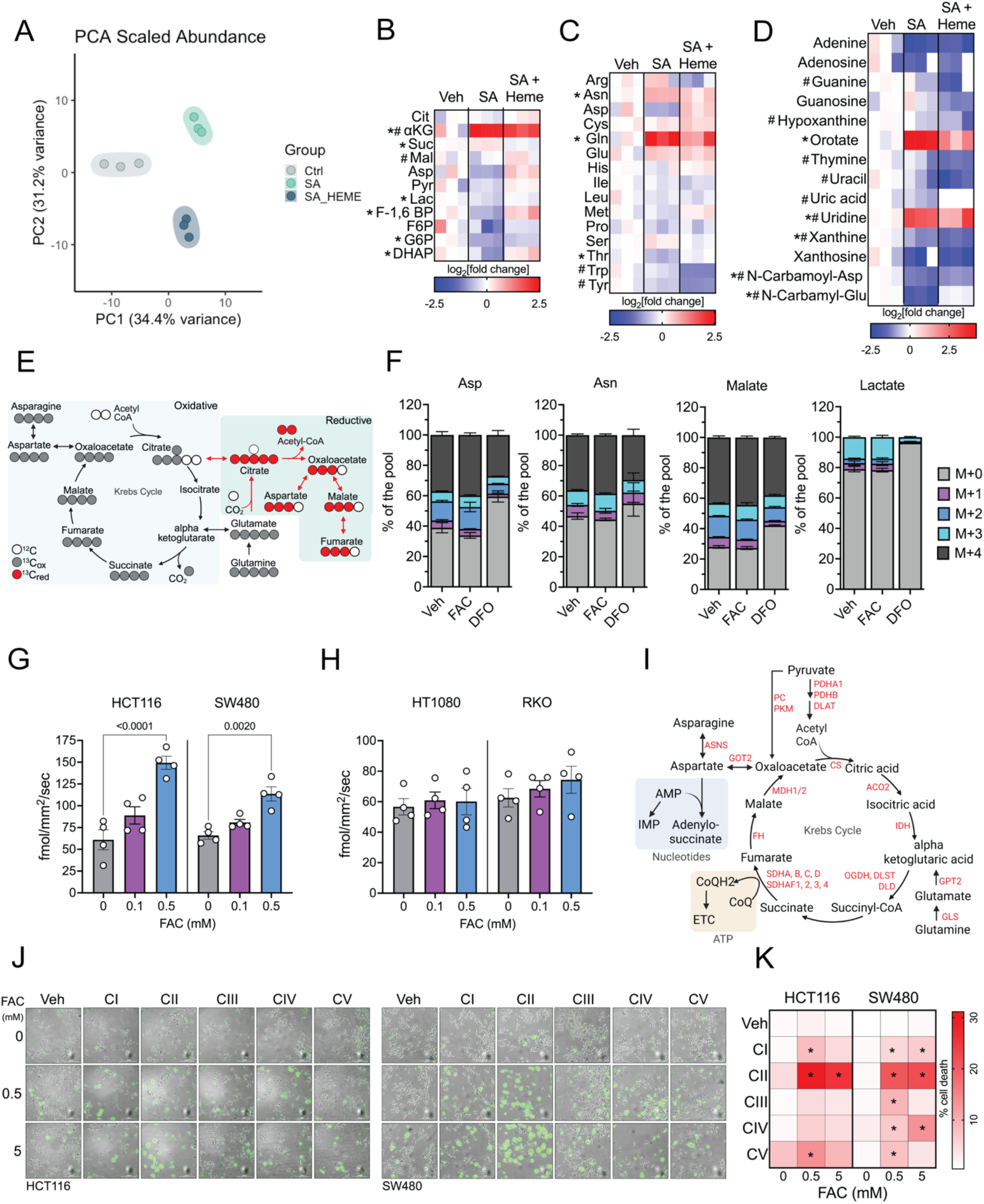
Mitochondrial complex II protects CRC cells from iron-induced oxidative cell death. A) Principal component analysis (PCA) plots of targeted metabolomics performed on HCT116 cells treated with FAC (250μM), SA (500μM) and heme (10μM) for 72h. B) Heatmap representation of log2 fold change in metabolites involved in TCA cycle, C) amino acids, D) metabolites de novo nucleotide biosynthesis, (* and # indicate p<0.05 between Veh and SA or # Veh and SA + Heme respectively). E) Schematic showing ^13^C_5_ L glutamine labeling strategy to trace glutamine consumption via TCA cycle to generate intermediates for de novo nucleotide synthesis pathway. F) % labeled asparagine, aspartate, malate, and lactate, 8h post 4mM 13C5 L-glutamine treatment. G) Oxygen consumption rate of HCT116, SW480 48h post FAC (0.1 and 0.5mM) or heme (10μM) treatment measured using Resipher^TM^ oxygen sensor, normalized with cell count measured by brightfield microscopy. H) Oxygen consumption rate of HT1080 and RKO under 48h post FAC (0.1 and 0.5mM) or heme (10μM) treatment measured using Resipher^TM^ oxygen sensor, normalized with cell count measured by brightfield microscopy. I) Schematic showing TCA cycle and its contribution to mitochondrial respiration. J) Representative Sytox green staining images of HCT116 and SW480 cells treated with FAC (0.5, 5mM) with or without inhibitors of the mitochondrial electron transport chain complexes (CI: Piericidin, 10nM, CII: TTFA, 100μM, CIII: Antimycin A, 100μM, CIV: KCN, 2.5mM, CV: Oligomycin, 1μM) for 24h K) Heatmap representation of % cell death calculated from J) * indicate p<0.05 as compared to Vehicle control +/- FAC. All data are mean ± SEM. One-way ANOVA with Tukey’s multiple comparisons test for (B), (C), (D), (G) and (K). ∗p < 0.05, ∗∗p < 0.01, ∗∗∗p < 0.001.

To probe whether elevated iron enhances flux through these pathways, we performed ^13^C₅-L-glutamine tracing (Fig. 4E). FAC treatment did not significantly increase incorporation of label into metabolites altered by heme deficiency, whereas iron chelation with deferoxamine reduced flux (Fig. 4F). Nevertheless, both FAC and heme supplementation increased OCR in iron resistant (HCT116 and SW480) but not in iron sensitive (HT1080 and RKO) cells, suggesting that the heme-dependent metabolic rewiring of the TCA cycle and nucleotide synthesis occurs primarily through respiratory chain–mediated processes (Fig. 4G-I). Given the observed ETC defects upon heme depletion, we hypothesized that disruption of specific complexes may sensitize CRC cells to iron stress. Indeed, pharmacologic inhibition of succinate dehydrogenase (SDH) selectively sensitized CRC cells to iron-induced toxicity, while inhibition of other ETC complexes produced modest effects (Fig. 4J, K). These findings point to the heme–CII axis as a central metabolic safeguard that enables CRC cells to tolerate high iron and sustain proliferation.

### Succinate dehydrogenase (SDH)–CoQ axis integrates mitochondrial metabolism with iron detoxification

CII is composed of four nuclear-encoded SDH subunits that mediate electron transfer from the TCA cycle into the ETC through succinate oxidation to fumarate. Electrons released from this reaction pass sequentially through FAD, three iron–sulfur clusters, and a heme cofactor before ultimately reducing CoQ, which then serves as an electron carrier to CIII^36^(Fig. 5A). Similar to pharmacological inhibition of CII by (2-Thenoyltrifluoroacetone) TTFA, CRC cells deficient in the structural subunits SDHA or SDHB exhibited heightened sensitivity to iron toxicity but not to other stressors such as copper, H₂O₂, or 5-fluorouracil (Fig. 5B-C; Fig S6A-E). Interestingly, inhibition of succinate oxidation within the TCA cycle using competitive inhibitors (malonate, succinate) did not alter iron sensitivity, whereas inhibition of CoQ reduction (via TTFA) did (Fig. S6F), suggesting that the CoQ-reducing function of CII, rather than its dehydrogenase activity, is critical for iron homeostasis.

**Figure 5:**
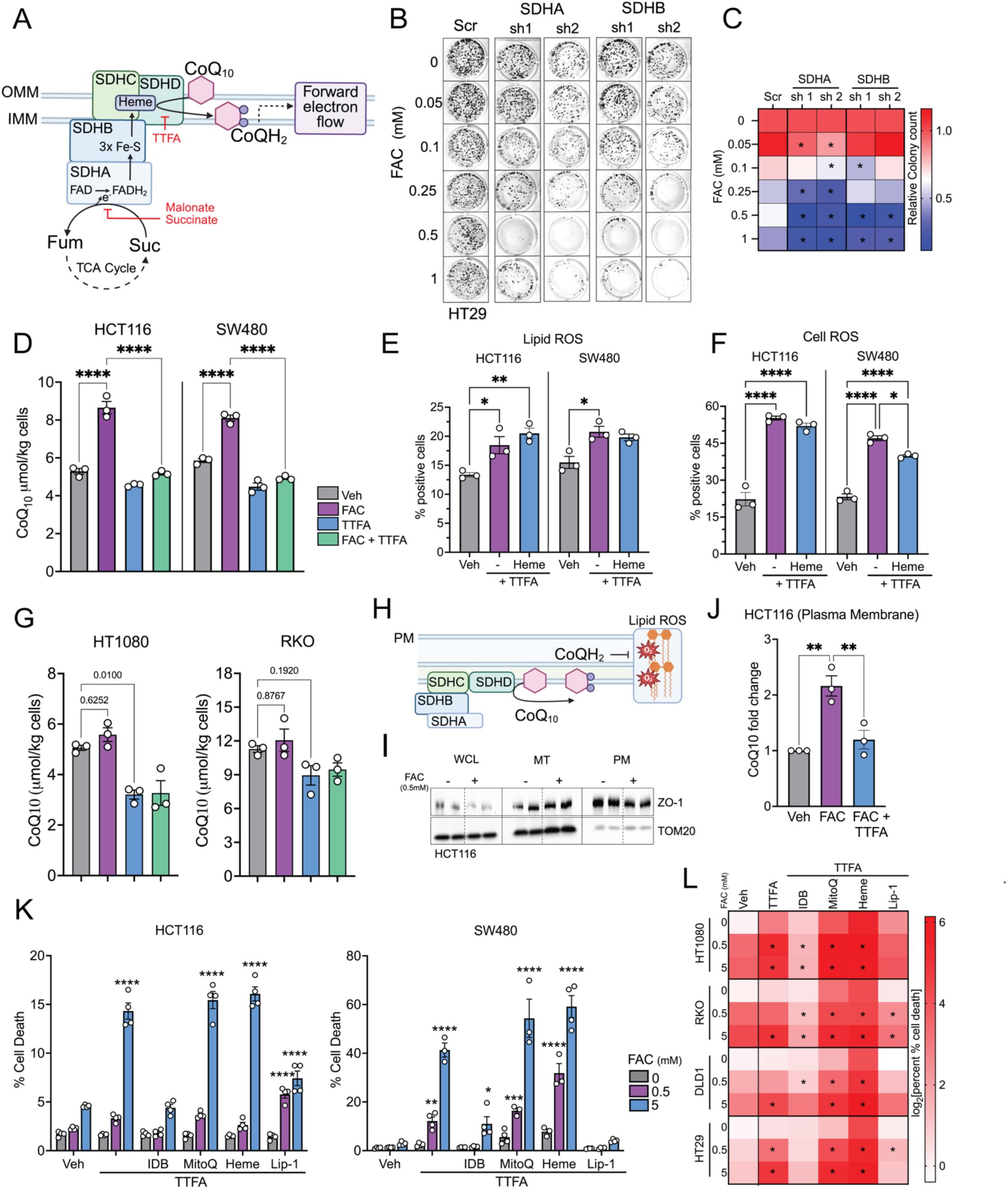
Succinate dehydrogenase (SDH)–CoQ axis integrates mitochondrial metabolism with iron detoxification. A) Schematic showing flow of electrons generated from succinate oxidation to through the different subunits of succinate dehydrogenase (CII) and ultimately reduce CoQ10 a terminal e-acceptor. B) Representative crystal violet staining images of HT29 cells stable for empty pLKO-puro (Scr) or shRNA targeting SDHA or SDHB (shRNA 1 or shRNA 2) 14 days post FAC treatment. C) Heatmap representation of relative colony count measured from B), * indicate p<0.05 compared to Vehicle treated control group. D) Total CoQ_10_ levels in HCT116 and SW480 cells treated with FAC (500μM) for 72h, with or without TTFA (100μM) for 24h measured by HPLC. E) Lipid ROS measured by C11-BODIPY581/591 in HCT116 and SW480 cells 24h post TTFA (100μM) and Heme (10μM) treatment. F) General ROS measured by free radical sensor carboxy-H2DCFDA in HCT116 and SW480 cells 24h post TTFA (100μM) and Heme (10μM) treatment. G) Total CoQ_10_ levels in HT1080 and RKO cells treated with FAC (500μM) for 72h, with or without TTFA (100μM) for 24h measured by HPLC. H) Schematic showing role of CoQ_10_ in mitigating lipid ROS. I) Western blot showing expression of ZO-1 and TOM20 as plasma membrane and mitochondria specific proteins. J) Relative CoQ_10_ levels in the plasma membrane (PM) in HCT116 cells treated with 0.5mM FAC for 72h using HPLC. K) % cell death measured by Sytox green staining in HCT116 and SW480 cells treated with FAC (0.5, 5mM) and TTFA (100μM) with or without heme (10μM) and CoQ analogs (Idebenone: IDB, 4μM, MitoQ: 20mM, Liproxstatin: Lip-1, 2μM) for 24h L) Heatmap representation of log2 fold change in % cell death in HT1080 (non-CRC) and RKO, DLD1, HT29 (CRC) cell lines treated as described in (K), * indicate p<0.05 compared to Vehicle treated control group. All data are mean ± SEM. One-way ANOVA with Tukey’s multiple comparisons test for (C), (D), (E), (F), (G), (J), (K) and (L). ∗p < 0.05, ∗∗p < 0.01, ∗∗∗p < 0.001.

We next considered whether heme availability, which regulates multiple ETC complexes (Fig. 4H-I), could influence CII activity^37^. Heme depletion did not significantly alter cellular succinate levels; however, we found that CII-dependent CoQ reduction was essential for buffering iron toxicity (Fig. 5D). Indeed, both total and reduced CoQ levels increased in cells exposed to elevated iron in a CII-dependent manner. Inhibition of CII-dependent CoQ reduction increased both lipid ROS and total ROS, effects that could not be rescued by heme supplementation (Fig. 5E-F). Interestingly increasing intracellular iron via FAC treatment did not induce total CoQ_10_ levels in iron sensitive HT1080 and RKO cells (Fig 5G).

Given prior data that CoQ can translocate from mitochondria to other cellular membranes, where it contributes to detoxification of lipid ROS, we examined CoQ distribution under iron stress(Fig 5H)^38,39^. Remarkably, we observed an iron-induced, CII-dependent increase in plasma membrane CoQ levels, further supporting its role as a lipid antioxidant defense system (Fig. 5I-J). In contrast, deletion of CI subunits (NDUFS1, NDUFS2) did not alter iron sensitivity, underscoring the specificity of the heme–SDH–CoQ axis in protecting against iron toxicity (Fig. S6G). Notably, CII inhibition–induced iron sensitivity was rescued by exogenous CoQ analogs and the ferroptosis inhibitor liproxstatin-1, but only the membrane-permeable CoQ analog idebenone—not the mitochondria-targeted MitoQ—restored viability (Fig. 5J–L). Together, these findings identify succinate dehydrogenase–dependent CoQ reduction as a central metabolic defense against iron-induced oxidative stress, linking TCA cycle electron flow, heme metabolism, and extramitochondrial antioxidant capacity across mitochondrial and plasma membrane compartments.

### Heme dependent SDHC expression is essential for protection against iron induced toxicity in CRC cells

Heme depletion led to a loss of multiple ETC subunits, diminished respiratory output, and a profound shift in the cellular metabolome. In erythroid progenitors, heme availability is known to transcriptionally and translationally control essential processes such as proliferation, differentiation, and intercellular communication^40^. However, whether similar heme-dependent regulation operates in cancer cells has remained unclear. To address this, we performed quantitative proteomics using tandem mass tag labeling of mitochondrial-enriched fractions from HCT116 cells treated with FAC, SA, or their combination for 72 h (Fig. S7A, B; Fig. 6A, B)^41^. Among the most striking changes, CII subunits and assembly factors emerged as iron-induced, heme-dependent proteins. SDHC (succinate dehydrogenase subunit C) was significantly enriched under FAC (heme sufficiency) but strongly depleted with FAC + SA (Fig. 6C). Heme-dependent SDHC stabilization was required to sustain CII activity and maintain cellular CoQ pools, thereby mitigating iron toxicity in CRC cells. Consistently, heme depletion and CII inhibition yielded highly concordant proteomic signatures, with ∼48% overlap in differentially altered proteins (Fig. 6D, E). Notably, heme biosynthetic enzymes were largely unaffected by SA treatment, with the exception of ALAS1, which is regulated by cellular iron via IREs (Fig. S7C)^42^. Beyond CII, protein abundance across CI–CV subunits and iron–sulfur cluster–containing proteins was increased under FAC and reduced with SA, whereas Fe–S biosynthetic enzymes remained unchanged (Fig. S7D–F). Importantly, other CoQ-reducing enzymes that have also been implicated in mediating mitochondrial redox balance (DHODH, SQOR, ETFDH)^43,44^ were upregulated in FAC + SA cells, underscoring the central role of reduced CoQ as an antioxidant defense (Fig. S7G).

**Figure 6:**
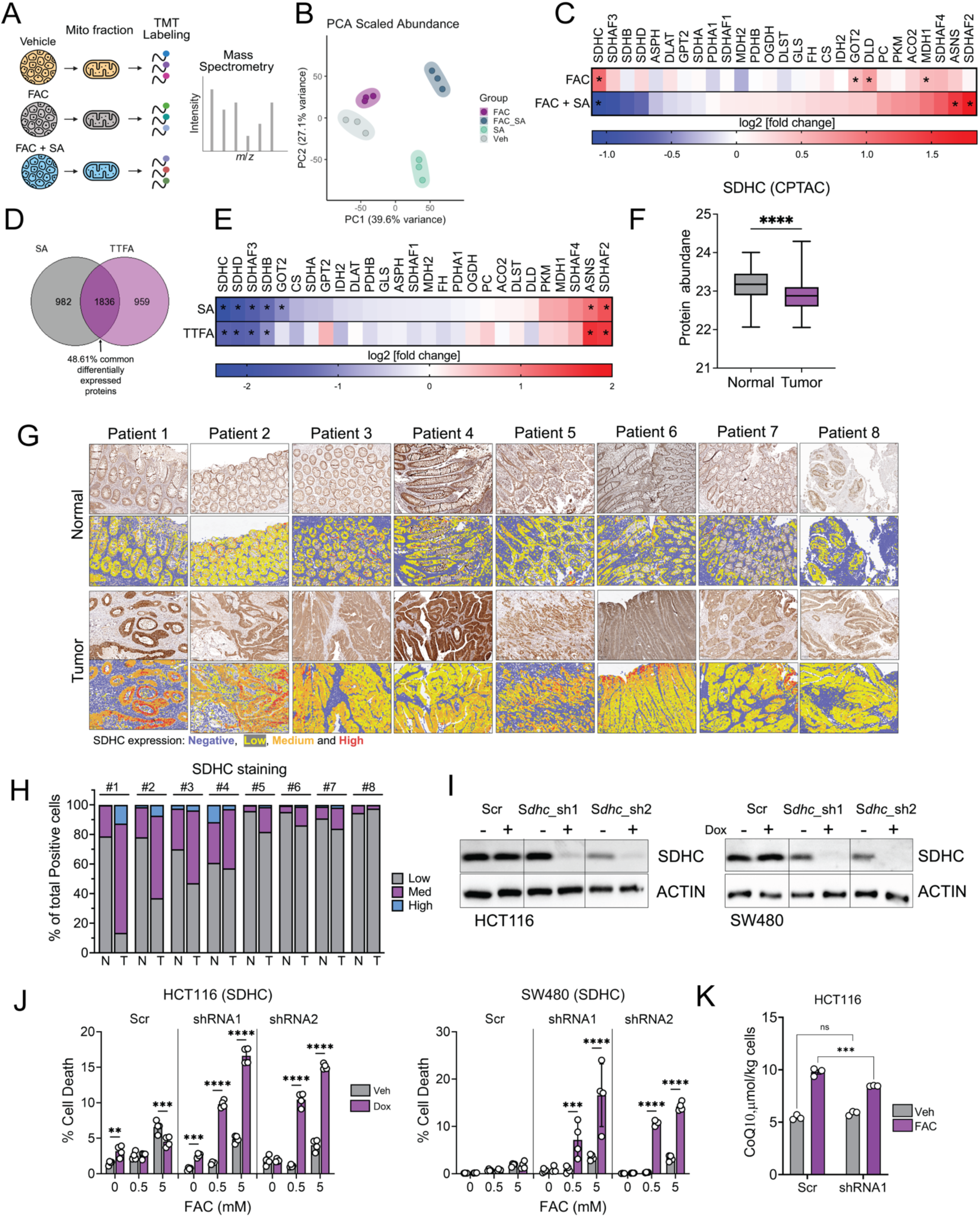
Heme Dependent SDHC Expression is Essential for Protection Against Iron Induced Toxicity in CRC Cells. A) Schematic showing the workflow for quantitative TMT labeling based proteomics in the mitochondrial enriched fractions of HCT116 cells 72h post FAC (500μM), SA (500μM) and Heme (10μM). B) Principal component analysis (PCA) plots of proteomics performed as described in A). C) Heatmap representation of log2 abundance fold change in cells treated with FAC and FAC + SA as described in A) D) Venn diagram showing the overlap between significantly altered proteins (p<0.05) upon treatment with SA, 500μM for 72h or TTFA, 100μM for 24h E) Heatmap representation of log2 abundance fold change in cells treated with SA and TTFA as described in D) F) Protein abundance of SDHC in Normal and tumor tissue of patients from CPTAC database. G) Representative Immunohistochemistry images showing expression of SDHC in normal and tumor tissue from patient tissue sections. The annotated images generated in Qupath categorized cells based on cell mean DAB OD and are shown as SDHC negative cells in blue and low, medium, high expression cells in yellow, orange, and red respectively). H) % low, medium, and high SDHC expressing cells represented as average of numbers computed for 3-5 different cross sections for each patient tissue section using Qupath as described in H) I) Western blot image showing SDHC expression and beta actin in HCT116 and SW480 cells stable for empty tet-pLKO-puro (Scr) or shRNA targeting SDHC (shRNA 1 or shRNA 2) 48h post Doxycycline treatment (250ng/mL). J) % cell death measured via Sytox green staining in HCT16 and SW480 cells (described in G), 48h post dox and FAC (0.5, 5mM) treatment. K) Total CoQ10 levels in HCT116 Scr or SDHC targeting shRNA1 cells 48h post dox or FAC (0.5mM) treatment measured by HPLC. All data are mean ± SEM. t test for (C), (E) and (F) or One-way ANOVA with Tukey’s multiple comparisons test for (J) and (K). ∗p < 0.05, ∗∗p < 0.01, ∗∗∗p < 0.001.

Interestingly, our data suggest that iron itself directly increases SDHC protein abundance in CRC cells, however, large-scale patient datasets demonstrate the opposite. Analysis of The Cancer Genome Atlas and Clinical Proteomic Tumor Analysis Consortium revealed that all SDH subunits (*SDHA–D*) are decreased at the mRNA and proteins level in CRC tumors relative to matched normal tissue (Fig. 6F)^45^. Our immunohistochemistry analyses demonstrated a clear increase in SDHC protein specifically within tumor epithelial cells of COAD patients as compared to the normal adjacent tissue (Fig. 6G, H). These findings demonstrate that bulk tumor profiling underestimates tumor intrinsic SDH expression as invasive CRC lesions are intermixed with surrounding stromal tissue, which expresses negligible SDH. Functional validation confirmed the tumor-specific requirement for SDHC. Inducible knockdown of *SDHC* in HCT116 and SW480 cells enhanced iron toxicity, phenocopying pharmacological inhibition of CII with TTFA, and concomitantly reduced iron-induced CoQ_10_ levels (Fig. 6I–K). Together, these findings define a heme-dependent regulatory mechanism that sustains SDHC expression and CoQ_10_ homeostasis, positioning the heme–SDH–CoQ axis as a critical antioxidant pathway that enables CRC cells to thrive under iron-rich conditions.

### SDHC inhibition sensitizes CRC cells to iron induced toxicity and reduce tumor progression

To evaluate the in vivo efficacy of SDHC inhibition in promoting iron-induced toxicity, we generated CT26 mouse CRC cells with a doxycycline-inducible shRNA targeting *Sdhc* and implanted them subcutaneously into Balb/c mice (Fig. 7A; Fig. S8A). Once tumors were established, doxycycline-containing chow was administered to induce *Sdhc* knockdown. SDHC-deficient tumors were significantly smaller and exhibited elevated lipid ROS, although the magnitude of this effect was moderate (Fig. 7B–C; Fig. S8B-C). Because subcutaneous tumor models do not fully recapitulate the colonic tumor microenvironment or the extent of iron accumulation observed in patients, we next crossed *Sdhc*^F/F^ mice with a tamoxifen-inducible intestinal epithelial Cre line (Villin-ER^T2^Cre mice) to generate intestine-specific *Sdhc*-deficient mice (Sdhc-IntKO) (Fig. 7D)^9,46^. Immunoblot analysis confirmed near-complete loss of SDHC in *Sdhc*-IntKO mice, whereas mice retaining one wild-type allele (*Sdhc*-IntHet) showed an ∼50% reduction in *Sdhc* levels with no histological changes in the intestinal epithelium (Fig. 7E; Fig. S8D). Similar to *in vitro* findings, *Sdhc* deficient mice showed signs of GI toxicity when fed high 1000ppm iron containing diet (Fig. 7F-G). *Sdhc* deficient mice on high iron diet for one week, lost up to 25% of total body weight with significant damage to the epithelial lining and accumulation of lipid peroxides as assessed by 4-hydroxynonenal staining (Fig. S8E-G; Fig. 7H). Because complete *Sdhc* loss in the intestine is predicted to compromise epithelial viability and confound tumorigenesis studies, we focused on the heterozygous model to test whether partial loss of *Sdhc* is sufficient to sensitize tumors to iron-induced stress^46^.

**Figure 7:**
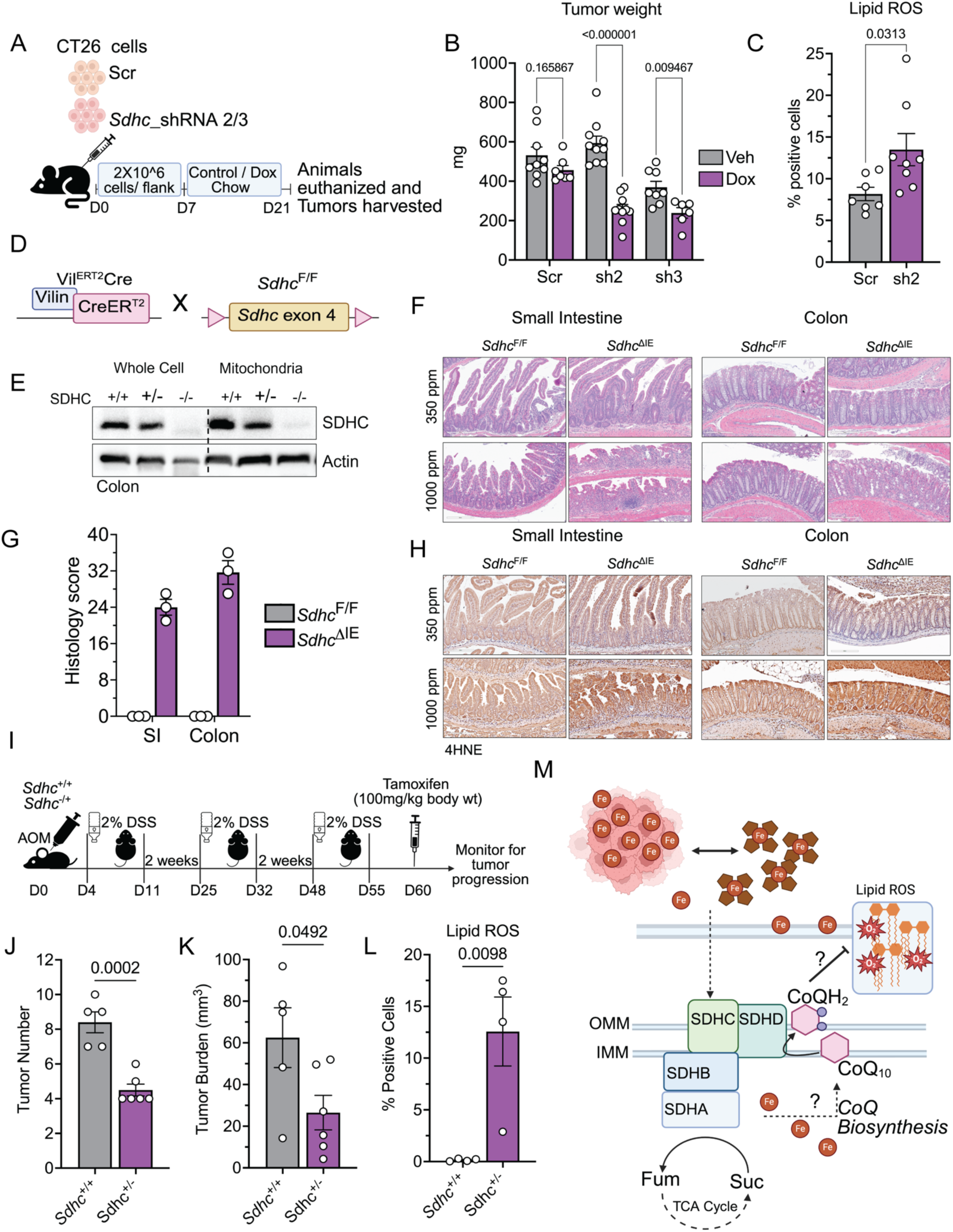
SDHC Inhibition Sensitizes CRC Cells To Iron Induced Toxicity and Reduce Tumor Progression. A) Schematic showing the xenograft tumor model with subcutaneous implantation. B) Tumor weight from B) C) Lipid ROS are measured by C11-BODIPY581/591 in EPCAM positive cells from the tumors from B). D) Schematic showing breeding strategy to generate tamoxifen inducible intestine specific SDHC deficient mice. E) Western blot images showing cellular and mitochondrial protein expression of SDHC 5days post tamoxifen induction via 100mg/kg body weight I.P. injection in WT (*Sdhc*+/+), Het (*Sdhc*+/-) and KO (*Sdhc*-/-) mice. F) Representative H&E images of *Sdhc*^F/F^ and *Sdhc*^ΔIE^ mice fed with 350 ppm or 1000ppm iron containing chow for 7 days. G) Inflammatory histology scored for small intestine and colon of *Sdhc*^F/F^ and *Sdhc*^ΔIE^ mice fed with 350 ppm or 1000ppm iron containing chow for 7 days. H) Representative immunohistochemistry images showing accumulation of 4 hydroxynonenal (4HNE) in small intestine and colon of *Sdhc*^F/F^ and *Sdhc*^ΔIE^ mice fed with 350 ppm or 1000ppm iron containing chow for 7 days. I) Schematic showing treatment protocol for colitis associated cancer model using AOM/DSS in *Sdhc*^F/F^ (WT) and *Sdhc*^Het^ ^IE^ (Het) mice. J) Tumor number measured in mice from I). K) Tumor burden measured in mice from I) L) Lipid ROS measured by C11-BODIPY581/591 in EPCAM positive epithelial cells derived from I). M) Schematic showing role of heme-SDHC-CoQ axis in mitigating iron induced cell death in CRC cells. All data are mean ± SEM. One-way ANOVA with Tukey’s multiple comparisons test for (B) and (G) and t test for (C), (J), (K) and (L). ∗p < 0.05, ∗∗p < 0.01, ∗∗∗p < 0.001.

To this end, mice were treated with azoxymethane (AOM) and subjected to three cycles of DSS to induce colitis-associated cancer. After completion of the three DSS cycles, at a stage when tumors are reliably established, tamoxifen was administered to delete one copy of *Sdhc* (Fig. 7I). Strikingly, *Sdhc*-IntHet mice developed markedly fewer tumors than their littermate controls, accompanied by increased lipid ROS and overall ROS accumulation (Fig. 7J-L; Fig. S8H-J). Collectively, these findings define a previously unrecognized heme–SDHC axis that regulates iron-induced toxicity in vivo models (Fig. 7M).

## Discussion

CRC cells are highly dependent on iron to support proliferation by fueling nucleotide synthesis and mitochondrial respiration. Yet, excess iron generates ROS and lipid peroxidation, which can trigger ferroptotic cell death. How CRCs thrive under these iron-rich conditions, which are cytotoxic to most other cell types, has remained poorly understood. While prior studies have largely focused on iron import, storage, and export via transferrin receptor, ferroportin, and ferritin, little is known about the cellular mechanisms that actively buffer high intracellular iron^5,8^. Our study identifies a previously unrecognized heme–SDH–CoQ axis that mediates this protective function. Cellular iron availability sustains SDHC protein abundance which in turn supports CoQ biosynthesis, which together contribute to lipophilic antioxidant defense, neutralizing iron-induced ROS and enabling CRC cells to exploit iron-rich environments for growth. This mechanism reveals a fundamental strategy by which CRCs resolve the iron paradox and maintain proliferation under potentially cytotoxic conditions.

Iron dependent lipid ROS–driven cell death, or ferroptosis, has been extensively characterized in vitro. Canonical regulators such as SLC7A11 and GPX4 were largely discovered through chemical genetic screens and through synthetic compounds^47,48^. However, whether these pathways explain iron toxicity in physiological contexts remains unclear. In CRC, although canonical ferroptosis regulators are transcriptionally upregulated, their expression does not correlate with patient survival, and deletion of these genes fails to sensitize cells to iron or alter tumor progression. This suggests that alternative, physiologically relevant protective mechanisms exist. Several intrinsic (oncogenic mutations in KRAS, Tp53) and extrinsic (nutrient and oxygen availability) factors affect ferroptosis sensitivity in cancers; mechanistically these factors all converges to altered cellular iron homeostasis^49–52^. In PDAC, KRAS activation increases the labile iron pool via ferritinophagy to fuel mitochondrial metabolism via iron-sulfur cluster biogenesis, but also sensitizes cells to iron toxicity and consequently ferroptosis^53,54^. Interestingly, loss of iron sulfur clusters sensitize lung cancer cells to ferroptosis via IRP stabilization which increases the labile iron pool^55,56^. Our work positions SDH and CoQ as critical iron-specific antioxidant defenses in CRC cells, functioning in parallel to canonical ferroptosis pathways. By demonstrating that mitochondrial CoQ metabolism directly mitigates iron-induced lipid ROS, we expand the conceptual framework of ferroptosis and iron toxicity, particularly in vivo and in intestinal tissues where canonical ferroptosis effectors are dispensable.

SDH has classically been studied as a metabolic hub, integrating TCA cycle flux with ETC activity, and as a tumor suppressor when mutated in paragangliomas, GIST, and renal cancers^36,57–59^. It remains unclear whether these tumor types experience significant iron accumulation; if not, SDH mutations in this context may have limited impact on iron buffering. In such cases, the signaling and metabolic advantages conferred by SDH loss may outweigh any detrimental effects of decreased protection against iron-induced stress. In contrast, our work identifies a previously unrecognized role for SDH in safeguarding cells against iron toxicity. Specifically, heme-dependent maintenance of SDHC preserves CoQ pools that neutralize lipid ROS. Notably, SDH also serves as a key regulator of CoQ redox status through its bidirectional catalytic capacity to oxidize succinate or reduce fumarate^60,61^. This property underlies tissue-specific variation in SDH function. In some tissues such as liver, kidney, and brain constitutively engage more in fumarate reduction than succinate oxidation, suggesting that cancers arising in these contexts may be less protected against iron overload by complex II activity^62^. Indeed, both liver and kidney are highly susceptible to iron-induced oxidative damage and ferroptosis, consistent with the notion that limited SDH-dependent succinate oxidation may compromise CoQ redox buffering capacity in these tissues^63–65^. Inhibition of SDHC dependent succinate ubiquinone reductase (SQR) activity has also been shown to increase sensitivity to BCL-2 antagonist venetoclax in multiple myeloma while heme supplementation induces resistance^66,67^. Whereas targeting SDHA dependent succinate oxidation reduces de novo purine synthesis, sensitizing cancer cells to inhibition of purine salvage pathways^68^. These findings also highlight the multifaceted role of mitochondrial ETC complexes where specific catalytic activity of the same complex can have vastly different affects.

At present, the mechanisms by which iron or heme regulate SDHC remain unresolved. One possibility is that heme or iron regulates SDHC stability through mitochondrial quality-control pathways, which are tightly coupled to respiratory complex integrity^69^. Another intriguing possibility is that heme binding to SDHC or associated complex II subunits stabilize the holoenzyme, thereby protecting it from proteolytic degradation. How SDHC regulation leads to a net increase in the CoQ pool is also unclear, but it is possible that enhanced SDHC activity shifts electron flux or alters CoQ redox cycling, leading to increased CoQ synthesis, redistribution, or stabilization. Future work will be needed to understand these mechanisms.

Beyond its canonical role as an electron carrier in the ETC, CoQ functions as a glutathione-independent, membrane-specific antioxidant^70–72^. Its redistribution to non-mitochondrial membranes lead to protection from iron-induced lipid peroxidation^38,39^. CoQ has been described as an radical trapping antioxidant in the membrane where it is reduced by FSP-1 and DHODH ^43,70,71^. In this sense, the heme–SDH–CoQ axis may operate as a mitochondrial counterpart to the FSP1–CoQ system, with mitochondria actively supplying CoQ to the plasma membrane and other cellular compartments. This model is supported by the increased abundance of other mitochondrial CoQ reducing enzymes under heme deficiency where SDH is no longer functional. However, it remains unclear how CoQ is redistributed to non-mitochondrial membranes in response to iron. Our proteomic data did not detect changes in STARD7, a recently described CoQ transporter, suggesting that other factors may mediate this process^73^. These findings highlight a broader principle: cancer cells co-opt redox-active cofactors, such as heme and CoQ, not only to sustain metabolism but also to adapt to oxidative stress.

Collectively, these findings uncover a novel mitochondrial antioxidant defense mechanism, redefine the landscape of ferroptosis regulation in vivo, and illuminate how CRCs buffer iron toxicity. By delineating the heme–SDH–CoQ axis, our study reveals a previously unknown metabolic adaptation that underlies CRC survival in iron-rich environments and highlights potential therapeutic opportunities to exploit iron-dependent synthetic vulnerabilities. In particular, our data suggest that pharmacologic inhibition of CII could disrupt this protective axis. When combined with approaches that elevate labile iron or impair parallel antioxidant defenses, such strategies could offer a synergistic means of inducing synthetic lethality in cancer cells. Thus, identifying the heme–SDH–CoQ axis not only advances our understanding of iron biology in cancer but also provides a promising therapeutic in CRC.

## Experimental Models

### Mice

All mice used in this study are predominantly C557BL/6 background unless mentioned otherwise. 6-8 weeks old male and female mice were used for all the experiments and littermates were randomized in all experimental conditions. All the mice were housed at Unit for Laboratory Animal Management at University of Michigan in standard temperature controlled specific pathogen free environment with 12h light/dark cycle and had excess to standard chow and water ad libitum. Mouse lines used are Villin-ER^T2^Cre;*Gpx4*^F/F^ ^20^. To generate GPX4 deficient mice with *Apc*, *Tp53* and *Kras* mutation, *Gpx4*^F/F^ mice were crossed CDX2-ER^T2^Cre;*Apc*^F/F^ mice (single mutant), CDX2-ER^T2^Cre;*Apc*^F/F^,*Tp53*^F/F^ mice (double mutant-p53), CDX2-ER^T2^Cre;*Apc*^F/F^,*Kras*^G12D^ mice (double mutant-KRAS) and CDX2-ER^T2^Cre;*Apc*^F/F^, *Tp53*^F/F^,*Kras*^G12D^ mice (triple mutant)^21^. And finally to generate intestine specific SDHC deficient mice, Villin-ER^T2^Cre mice(Jax strain 032770) were crossed with *Sdhc*^F/F^ mice^46^. All animal studies were performed in accordance with the Association of Assessment and Accreditation of Laboratory Animal Care International guidelines, approved by the University Committee on the Use and Care of Animals, University of Michigan.

#### Tamoxifen induction

Mice were intraperitoneally administered with 100mg/kg body weight tamoxifen dissolved in corn oil every other day 3 times. Mice were used for experiments 5 days post last tamoxifen dose unless mentioned otherwise.

#### AOM-DSS induced colitis associated cancer (CAC) model

Mice were injected with 10mg/kg body weight azoxymethane (AOM) intraperitoneally on Day 0. Day 3 mice were cycled on and off with 2% dextran sodium sulfate (DSS) in drinking water for 7 days, while body weights were recorded every other day^20^. 5 days after the last DSS cycle, mice were induced with tamoxifen as described above and euthanized 4 weeks after last DSS cycle, tissues were either flash frozen, fixed and Swiss rolled or processed live for molecular measurements, histological analysis for ROS measurements respectively.

#### Syngeneic and Xenograft studies

Wild type BalbC mice were anesthetized with 2-2.5% isoflurane and 2 million CT26 cells stable (Scr, SDHC-sh2/3) were implanted into the lower flanks subcutaneously. Mice were then placed on standard or doxycycline containing chow 7 days after the implantation after the tumors were palpable. Mice were euthanized roughly 2 weeks after or the tumors reached 2cm^3^ in size, the humane end point^26^. Tumor volume/weight was measured, and the tissues were either flash frozen, fixed and Swiss rolled or processed live for molecular measurements, histological analysis for ROS measurements respectively.

#### High Iron Diet

For high iron studies mice were given tamoxifen to induce Cre-mediated recombination as described above. After administering the last tamoxifen dose, mice were fed with standard diet chow or high iron containing chow i.e., AIN-93G Purified Rodent Diet chow with 350 ppm Fe (Dyets, 115180) or 1000ppm Fe (Dyets, 115842) added from Ferric Citrate respectively for 7 days. Body weight was recorded every day and % body weight change was calculated i.e. (body weight on day 7/ body weight on day 1) *100.

### Human Subjects

Paraffin embedded blocks of de-identified human adult colon biopsies were collected with approval from the University of Michigan Institutional Review Board. 10uM sections were cut from the blocks and uses for Immunohistochemistry (IHC). Deidentified human CRC enteroids and matched control enteroids were established and cultured as previously described^20^.

#### Cell lines

Cancer cell lines HCT116, SW480, DLD1, KM12, RKO-1, HT29, MC38, CT26, CD-1, NIH3T3, HT1080, DANG, ASPC, UOS2 were obtained from ATCC and maintained at 37°C, 5% CO_2_, 21% O_2_ humidified environment in DMEM supplemented with 10% heat inactivated fetal bovine serum (FBS) and 1% antibiotic/antimycotic agent).

### Method Details

#### Clonogenic Assay

250-500 cells were seeded in a 12 well plate in biological triplicates with 1mL media. The cells were treated as described in the legends for Fig. 2C, D; Fig S2F; Fig. 3C, D; Fig. S3B, E, G; Fig. S4; Fig. S5B-D, F-G; Fig 5B; 24h. 2 weeks after the treatment, cells were fixed in 10% buffered formalin and stained with 0.5% crystal violet, 20% methanol solution. Colonies were then blindly counted using Cytation 5 Imaging Multi-Mode reader.

#### Cell Growth Assays

Cell growth was measured using live cell imaging by the Cytation 5 Imaging Multi-Mode reader with attached BioSpa. 1000 cells/well were seeded in 96 well clear flat bottom plates and treated 24h later in biological triplicates or quadruplicate as described in the legends for Fig. 2A-B; Fig. S2D-E; Fig. 3G; Fig. S4B-C; Fig. S6C. Images were taken every 24h and normalized to the baseline values counted from images taken right before the treatments (0h). Relative growth was plotted using Graphpad Prism software.

#### Cell Death Assay

Cell death was measured using SYTOX Green Nucleic Acid stain (20nM, Invitrogen #S7020). 3500-5000 cells / well were seeded in 96 well clear flat bottom plates and treated 24h later in biological triplicates or quadruplicate as described in the legends for Fig. 2D; Fig. S2B-C; Fig. 4J-K; Fig. 5I-J; Fig. 6J. Imaged were taken every 8h using Cytation 5 Imaging Multi-Mode reader with attached BioSpa and % cell death was calculated by dividing number SYTOX^+^ cells by total number of cells.

#### Histology

Colonic and tumor tissues were rolled and fixed in 10% buffered formalin overnight, dehydrated in 70% ethanol and embedded in paraffin. 5μM thick sections were then stained for H&E and mounted with Permount Mounting Medium (Thermo Fisher Scientific). The slides were imaged using Leica Aperio Slide Scanner and representative images are shown at 10x or 20x magnification. Histological scoring was performed evaluating inflammation severity (0–3), depth of injury (0–3), and crypt damage (0–4). Each parameter was multiplied by the percentage of tissue involvement (1, 0–25%; 2, 26–50%; 3, 51–75%; 4, 76–100%), and the values were summed to yield a total injury score (maximum 40).

#### SDHC and 4HNE staining and quantification

For SDHC staining, 5μM thick sections obtained from the paraffin blocks containing human CRC tissues were rehydrated and antigen retrieval was done using 10mM sodium citrate buffer. Followed by blocking with 5% goat serum in PBS and sections were then probed with primary antibody against SDHC (Proteintech, 14575-AP) or 4HNE (R&D Systems, MAB3249-SP) diluted at 1:500 in the blocking solution. After 3 washes with PBS, slides were incubated HRP-conjugated anti-rabbit IgG at 1:500 dilution in the blocking solution for 1h. Sections were washed with PBS and incubated with DAB substrate and monitored for the color reaction (∼5minutes). The sections were then counter staining with hematoxylin, dehydrated and mounted with Permount mounting medium. The slides were imaged using Leica Aperio Slide Scanner and representative images are shown at 10x or 20x magnification. Qupath software was used for quantification and cells were categorized as SDHC negative, low, medium, or high based on mean cell DAB OD and represented as % of the total positive cells in Fig. 6H. The annotated images show SDHC negative, low, medium, and high cells as blue, yellow, orange, and red respectively (Fig 6. G).

#### Western Blotting

Protein lysates were prepared by lysing the cells or tissues in RIPA buffer supplemented with protease and phosphatase inhibitors. After lysis solubilized proteins were resolved on 12% SDS-polyacrylamide gels and transferred to PVDF membrane, blocked with 5% milk in TBST and probed overnight with indicated primary antibodies diluted at 1:1000 in the blocking buffer for GPX4 (Proteintech, 67763-1-Ig), FTH1 (Cell Signaling,3998S), UROD (Santa Cruz, SC-365297), FECH (Proteintech, 14466-1-AP), SDHA (Cell Signaling, 11998), SDHB (Proteintech, 10620-1-AP), SDHC (Proteintech, 14575-AP), β-actin (Proteintech 66009-1-Ig). Protein bands were then visualized by electrochemical luminescence using HRP conjugated secondary antibody against mouse (Cell Signaling, 7076S) or rabbit (Cell Signaling 7074S).

#### Real Time Quantitative PCR

RNA was isolated from cells or tissues using TRIzol chloroform method. RNA was reverse transcribed using MMLV reverse transcriptase using primers listed in the table 1. Quantitative PCR was done with Radiant Green qPCR Master Mix (Alkali Scientific Inc.) and primers listed in table 1. Quantification cycle values were normalized to β-actin and expressed and log_2_ fold change.

#### CRISPR screen and Analysis

The human CRISPR metabolic gene knockout library was a gift from David Sabatini containing 29,790 sgRNAs targeting 2981 human metabolic genes. A total of 75 × 10^6^ cells were seeded at 0.5X10^6^ cells/mL in 6 well plates and transfected with the CRISPR library virus through spinfection at 1200g for 1h in presence of 10μM polybrene. 24h after spinfection the virus was removed by changing the media and after additional 24h, 1μM/mL puromycin was added transfected cells were selected for 3 days. About 30 million cells were pelleted and saved as doubling 0 population for normalization purposes described later. 30 million cells/ condition were then treated with 2.5mM FAC, 100μM FG4592 or vehicle control and passaged every 3-4 days until 14 doublings. Genomic DNA was isolated from 15 million cells from doubling 0 control and doubling 14 FSC, FG4592 and vehicle treated cells using DNeasy kit (Qiagen). sgRNA inserts were PCR amplified using Takara Ex Taq polymerase, purified, and sequenced using NextSeq Illumina. The sequence reads were mapped to the sgRNA reference library and abundance of sgRNA was quantified (normalized to doubling 0 abundances). For gene level differential score the log2 fold change abundance between vehicle, FAC and FG4592 for all sgRNA targeting the same gene was averaged. Finally, the gene score for each condition was calculated as median log2 fold change in abundance between FAC or FG492 and vehicle treated population.

#### Generation of shRNA/sgRNA stable cell lines

shRNA targeting UROD, FECH and SDHC were cloned into pLKO.1-Tet on plasmid to generate doxycycline inducible knock down. While shRNA targeting SDHA and SDHB were cloned into pLKO.1 plasmid to generate constitutively active knockdown cells. Corresponding empty vectors were used as controls. shRNA sequences were sequence validated and packaged into lentivirus by University of Michigan vector core. Cells were transfected through spinfection at 900g for 1h and selected for 2 weeks in puromycin (2μM/mL). All clones were validated by measuring protein expression 24, 48 and 72h post doxycycline (250ng/mL) induction.

#### Cell ROS and Lipid ROS measurement

For cells, 1million cells/well were seeded in 6 well plates and 24h later were treated as indicated in the legends of Fig. 3F, and Fig. 5E-F for 72h and 24h respectively. Cells were harvested using PBS-EDTA (5mM) buffer, washed with HBSS and stained for lipid ROS or cell ROS with 5μM C11-BODIPY (Thermo Fisher, D3861) and 10μM cell permeable free radical sensor carboxy-H2DCFDA (Thermo Fisher, C10444) for 30minutes at 37°C respectively. Cells were pelleted, washed, and resuspended in FACS buffer (1x PBS, 2%BSA, 1mM EDTA) for analysis on Cytek aurora spectrum flow analyzer. Fluorescence intensity was measured in the FITC channel for a minimum of 20,000 cells, data analyzed using FlowJo software and values expressed as percentage of positive cells^26^.

For tumor and colon tissue: Middle region of the colon or tumors were collected from mice as indicated in the legend of Fig. 7E, K. The tissues were digested with 1mg/mL collagenase type IV dissolved in RPMI for 30 minutes at 37°C. Cell suspension was then filtered, collected, washed, and resuspended in FACS buffer (1x PBS, 2%BSA, 1mM EDTA). A single cell suspension was stained with 7-AAD (1:200; BD Pharmingen, catalog number 559925) and APC anti-mouse CD326 (EpCAM) antibody (1:200; BioLegend, catalog number 118214) for 30 min on ice. Subsequently, the single suspension was stained with 5μM C11-BODIPY (Thermo Fisher, D3861) and 10μM cell permeable free radical sensor carboxy-H2DCFDA (Thermo Fisher, C10444) for 30minutes at 37°C respectively. On Cytek aurora spectrum flow analyzer, forward and side scatter was used to identify single cells while 7AAD negative, APC positive cells were identified as live epithelial or tumor cells. Fluorescence intensity for C11-BODIPY or carboxy-H2DCFDA was measured in the FITC channel for previously identified live cell population^20^. The data was collected for a minimum of 20,000 cells, analyzed using the FlowJo software and values expressed as percentage of positive cells.

#### Metabolomics

HCT116 cells were seeded at 1 million cells/well seeding density and treated with FAC (100μM), Succinyl Acetone (500μM) and heme (10μM) for 72h, 24h post seeding. Metabolites were extracted in 80% methanol, lyophilized, and reconstituted in 50:50 methanol water mixture, normalized based on protein concentrations obtained from a well processed in parallel. Samples were then analyzed on Agilent 1290 Infinity II Bio LC ultra-high performance liquid chromatography system paired with an Agilent 6470 QQQ^74^. Data was processed and analyzed via MassHunter Quantitative Analysis v10.Metabolite abundance was median centered across samples and statistical significance was determined by one-way ANOVA with a significance threshold of 0.05.

#### ^13^C_5_ L-Glutamine Labeling

HCT116 cells were seeded at 1million cells/ well in a 6 well plate and treated 24h later with 0.5mM FAC or 150μM DFO for 24h. For stable isotope tracing, glutamine free DMEM (Thermo Fisher) was supplemented with 2mM ^13^C_5_-L glutamine (Sigma), 1mM pyruvate, 10% FBS, and 1% antibiotic/antimycotic solution. A parallel plate was treated similarly except it was supplemented with regular ^12^C_5_-L glutamine for normalization purposes. The cells were labeled for 8h and metabolites were extracted as described above. Samples were run on a Thermo Scientific IQX OrbiTrap LCMS system using a Waters Acquity UPLC BEH amide 1.7 um 2.1 × 100 mm column with a Waters UPLC BEH Amide 1.7 um VanGuard Pre-Column column of amide for high throughput acquisitions. Solvent A consists of 10 mM ammonium acetate in water with 10 mM Ammonia pH 9.2. Solvent B consists of acetonitrile. Pump Seal wash and autosampler wash is 50% isopropanol with 0.1% formic acid. The LC profile is 0.2 ml/min of 85% B at 0-0.5 min: 0.5-15 min, 15%B; 15-17 min, 15% B: 17.1 min, 85%B; 22 min, 85% B. The column compartment temperature is maintained at 30 °C and the autosampler is at 5 °C. The injection volume is 3 µl. Data was analyzed using Compound Discoverer v3.4.

#### Heme measurement

Total cellular heme was measured using fluorometric assay that measures protoporphyrin IX fluorescence upon release of free iron from heme as previously described^75^. 1X10^6^ cells were seeded in 6 well plate and 24h later treated with FAC, Heme and Succinyl acetone for 72h as described in the legend for Fig. 3B. Cells were washed with PBS, harvested, and resuspended in 250μL 20mM oxalic acid and incubated overnight at 4°C in dark. Next day, 250uL of supersaturated prewarmed 2M oxalic acid was added to the cells to achieve a final 1M oxalic acid concentration. The samples were then split in half and boiled at 100°C for 1h while the other half was kept at room temperature for background normalization. The samples were centrifugated at full speed for 5inutes at room temperature, 50uL of the supernatant was transferred to 96 well black wall, clear bottom plates and fluorescence was measured at 400nM excitation and 604nM emission. The fluorescence of unboiled sampled representing background was subtracted from the fluorescence of boiled sample and the heme concentration was calculated using a standard curve of hemin chloride as μM.

#### Oxygen Consumption Rate

Cells were seeded (5000 cells/ well) in a 96 clear flat bottom plate and treated 24h later as described in the legend of Fig. 3H and Fig. 4G-H. Plate was then interfaced with Resipher system by placing the Resipher sensors over the plate as per manufacturer’s instructions for live, continuous monitoring of oxygen consumption. OCR values computed by the Lucid Lab software were normalized to cell numbers.

#### Proteomics

Mitochondria were isolated using 3 million cells were seeded in a 15cm cell culture plate in 25mL media and treated with 0.5mM FAC, 0.5mM SA for 72h and 50μM TFFA for 24h in biological triplicates. Cells were washed, harvested in ice cold PBS and resuspended in homogenization buffer (containing 200mM sucrose, 10mM TrisCl pH7.5, 10mM EDTA and protease inhibitor cocktail). Cells were then homogenized with 30-40 strokes in the dounce homogenizer. Homogenate was the centrifuged twice at 1000g for 10minutes at 4°C to removed nuclear pellet and the mitochondrial pellet was obtained by centrifuging the supernatant at 9000g for 20 minutes at 4°C. The obtained mitochondrial pellet was washed with the homogenization buffer twice to remove any cellular debris and nuclear contaminants. 100ug of mitochondrial proteins were then digested into peptides with trypsin, labeled with tandem mass tag labeling kit (ThermoFisher) according to manufacturer’s instructions, analyzed on mass spectrometer by the University of Michigan proteomics core. The values are represented as log2 fold change in abundance compared vehicle treated group.

#### Co-enzymeQ_10_ measurement

For whole cell CoQ measurements, 1.5million cells were seeded in 10cm cell culture plates in 20mL media in biological triplicates, 24h later they were treated as indicated in the legends of Fig. 5D-H and Fig. 6K. Cells were then washed, harvested, and pelleted in ice cold PBS in pre-weighed tubes and wet weight of cell pellets was recorded. Cell pellets were then rapidly resuspended in n-propanol for CoQ extraction, where volume of n-propanol was normalized based on cell pellet weight. Samples were allowed to stand at room temperature and centrifuged at 12,000g for 5 minutes at 10°C to avoid CoQ precipitation and stored at -80°C. HPLC analysis for CoQ measurement was performed on a Hypersil ODS column (150 × 4.6mm, 3μm bead size, thermo Fischer) as described previously described^76^. The samples were run twice, first to measured oxidized CoQ, followed by oxidation of reduced CoQ by p-benzoquinone to measure total CoQ. Reduced CoQ was calculated by total CoQ – Oxidized CoQ.

For plasma membrane CoQ levels, 3 million cells were seeded in 15cm cell culture plate in 25mL media, treated 24h later and homogenized as described above for mitochondrial extraction. The remaining supernatant after pelleting mitochondrial content was centrifuges at 100,000g for 1h at 4°C to get plasma membrane pellet. Wet weight of the plasma membrane pellet was determined, samples were prepared and analyzed as described above.

---

**Figure S1.**
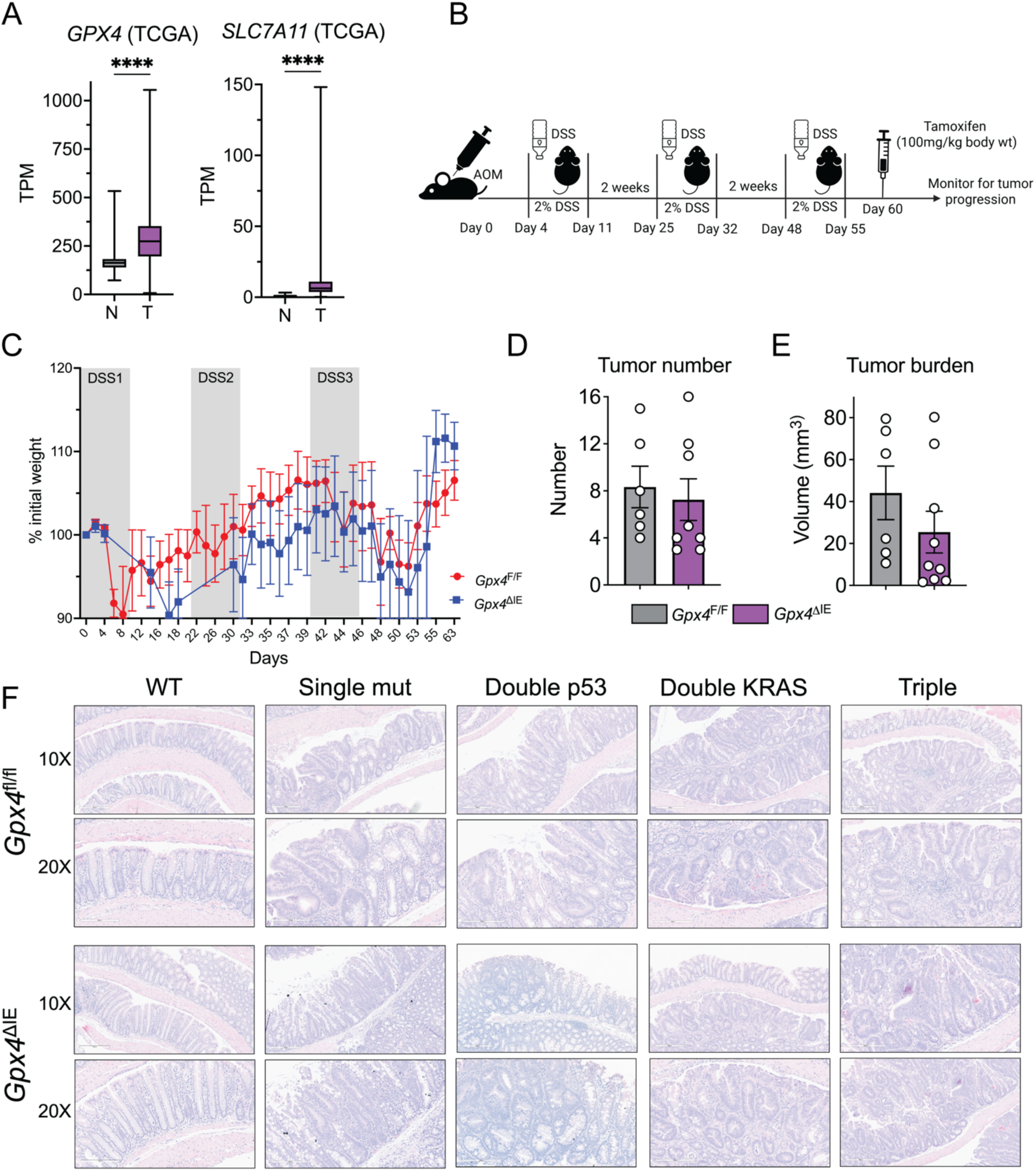
A) Gene expression of *GPX4* and *SLC7A11* in colon adenocarcinoma patients cancer and normal adjacent tissues from the TCGA cancer database. B) Schematic showing treatment protocol for colitis associated cancer model using AOM/DSS. C) % body weight change post AOM treatment in mice from (B) D) Tumor number from (B) E) Tumor burden from B) F) Representative H&E images of GPX4 deficient Single / Double-p53 / Double-KRAS^G12D^ / Triple mutant mice post tamoxifen induction via 100 mg/kg body weight I.P. injection showing regions of dysplasia in the colon. All data are mean ± SEM. t test for (A). ∗p < 0.05, ∗∗p < 0.01, ∗∗∗p < 0.001.

**Figure S2.**
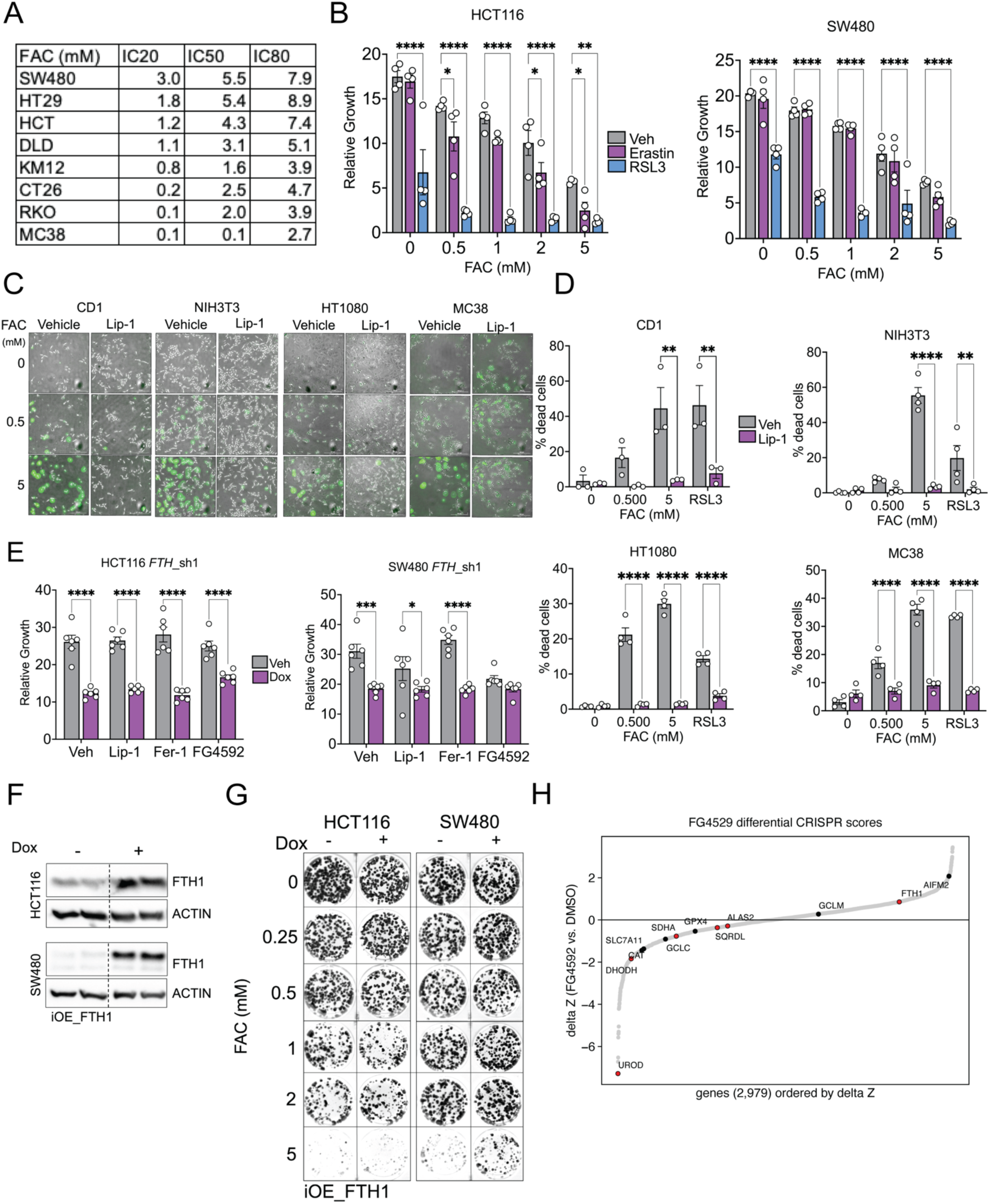
A) IC50, IC20 and IC80 values for human and mouse CRC cell lines treated with Ferric Ammonium Citrate (FAC) for 72h, calculated by measuring cell growth using bright field microscopy. B) Relative growth of HCT116 and SW480 cells 72h post treatment with FAC with or without ferroptosis inducers erastin (1μM) or RSL3 (1μM) using brightfield microscopy. C) Representative Sytox staining images from CD1, NIH3T3, HT1080 and MC38 cells 72h post FAC treatment with or without ferroptosis inhibitor Liproxstatin (Lip-1 2μM). D) % cell death quantification from (B). E) Relative growth of sh*FTH1* cells 72h post dox treatment with or without ferroptosis inhibitor liproxstatin (Lip-1 1μM), ferrostatin (Fer-1 1μM) or FG4592 (100μM) F) Western blot images showing protein expression of FTH1 and beta actin in HCT116 or SW480 cells stable of doxycycline inducible FTH1 overexpression construct 48h post dox treatment. G) Representative crystal violet staining images for cells from (F) 14 days post dox treatment with or without FAC treatment. H) Differential CRISPR score (DMSO-FG4592) highlighting significantly depleted genes of interest in doubling 14 populations. All data are mean ± SEM. One-way ANOVA with Tukey’s multiple comparisons test for (B), (D) and (E). ∗p < 0.05, ∗∗p < 0.01, ∗∗∗p < 0.001.

**Figure S3.**
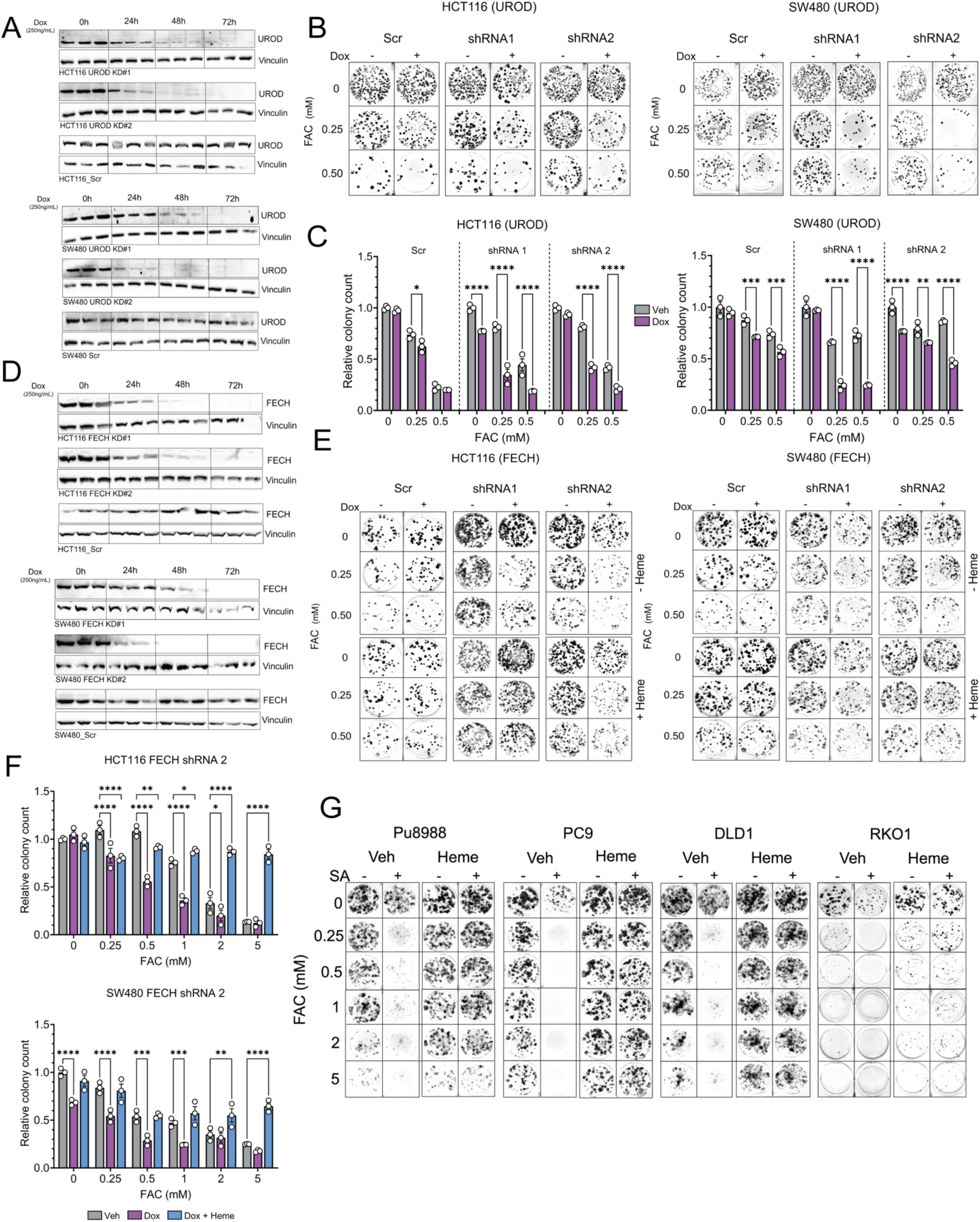
A) Western blot images showing protein expression of UROD and Vinculin in HCT116 and SW480 cells stable for empty tet-pLKO-puro (Scr) or UROD targeting shRNA (shRNA1 and shRNA2) 24h, 48h and 72h post dox treatment (250ng/mL). B) Representative crystal violet staining image in cells from b) 14 days post FAC treatment (0.25, 0.5mM) with or without dox (250ng/mL). C) Measurement of relative growth by quantification of relative colony count from (B). D) Western blot images showing protein expression of FECH and Vinculin in HCT116 and SW480 cells stable for empty tet-pLKO-puro (Scr) or FECH targeting shRNA (shRNA1 and shRNA2) 24h, 48h and 72h post dox treatment (250ng/mL). E) Representative crystal violet staining image in cells from e) 14 days post FAC treatment (0.25, 0.5mM) with or without dox (250ng/mL) or heme (10μM). F) Measurement of relative growth by quantification of relative colony count from (E). G) Representative crystal violet staining image in human CRC cells from 14 days post FAC treatment with or without SA (250μM) and Heme (10μM). All data are mean ± SEM. One-way ANOVA with Tukey’s multiple comparisons test for (C), and (F). ∗p < 0.05, ∗∗p < 0.01, ∗∗∗p < 0.001.

**Figure S4.**
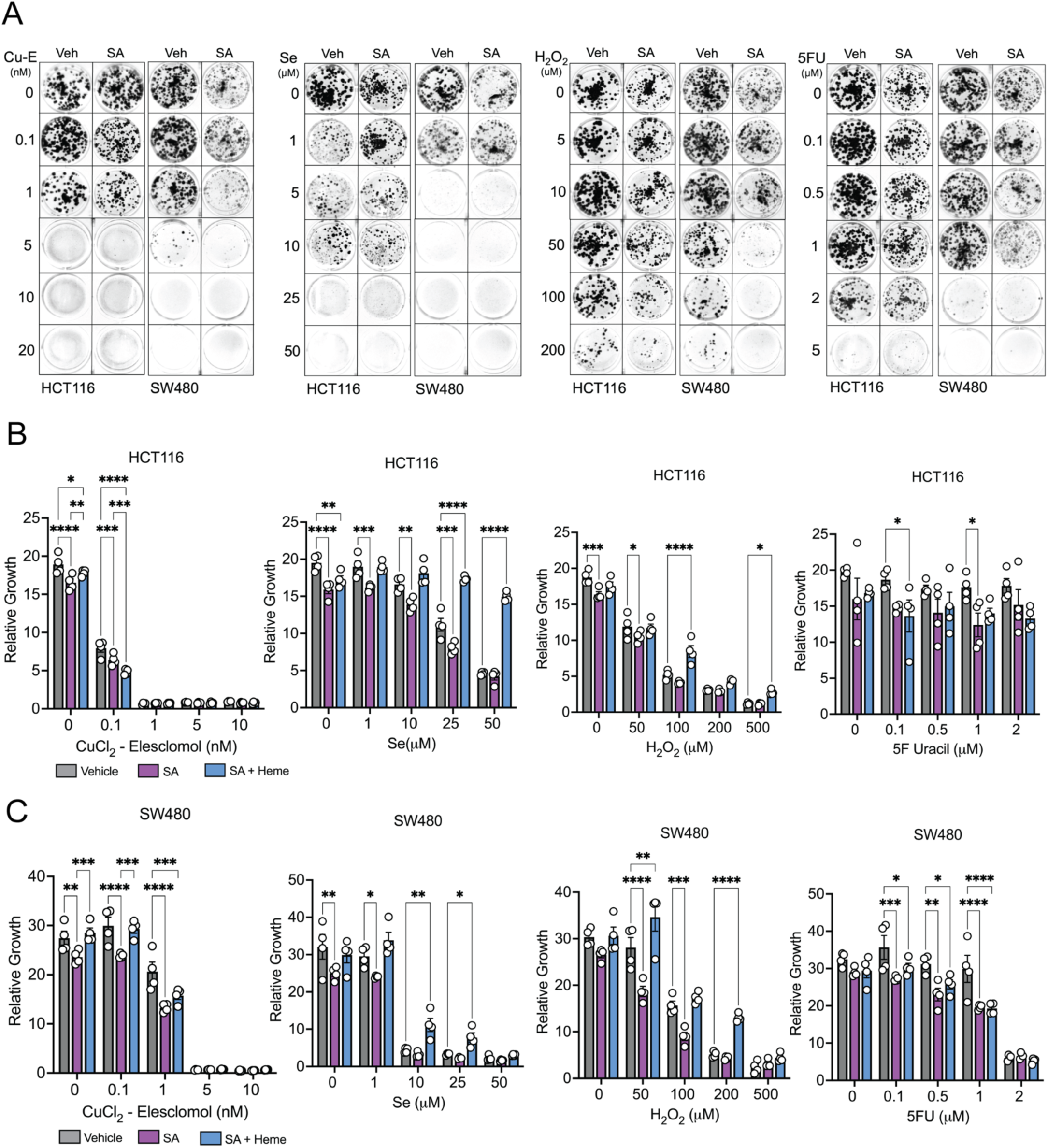
A) Representative crystal violet staining of HCT116 and SW480 cells 14 days post SA (250μM), copper sulphate (100μM) + elesclomol (Cu-Es), Sodium selenite (Se), hydrogen peroxide (H_2_O_2_) and 5-fluoroUracil (5FU) at indicated concentrations. B) Relative growth of HCT116 measured by bright field microscopy, 72h post SA (250μM), copper sulphate + elesclomol (Cu-Es), Sodium selenite (Se), hydrogen peroxide (H_2_O_2_) and 5-fluoroUracil (5FU) at indicated concentrations. C) Relative growth of SW480 measured by bright field microscopy, 72h post SA (250μM) copper sulphate + elesclomol (Cu-Es), Sodium selenite (Se), hydrogen peroxide (H_2_O_2_) and 5-fluoroUracil (5FU) at indicated concentrations. All data are mean ± SEM. One-way ANOVA with Tukey’s multiple comparisons test for (B), and (C). ∗p < 0.05, ∗∗p < 0.01, ∗∗∗p < 0.001.

**Figure S5.**
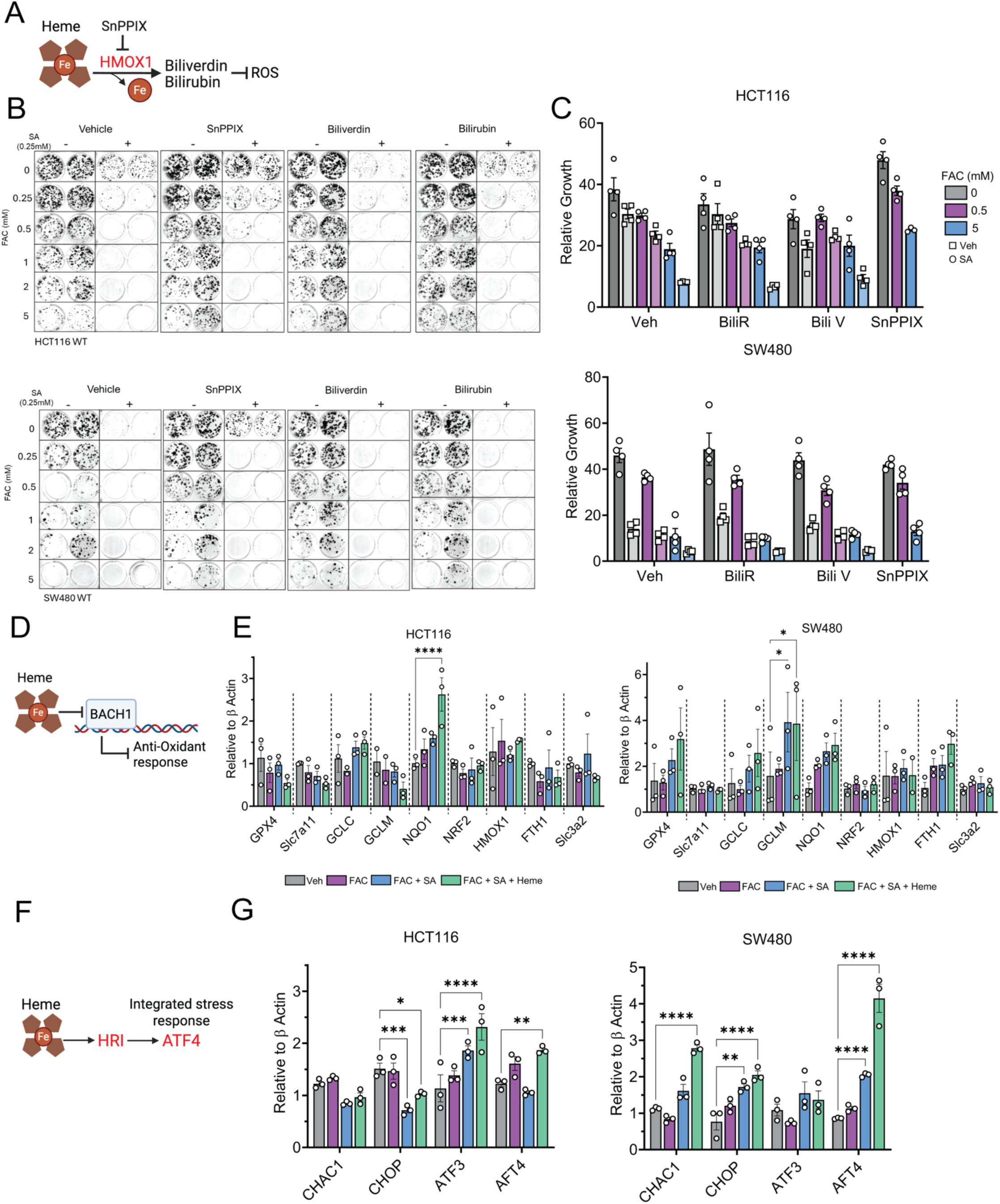
A) Schematic showing hemoxygenase (HO-1) dependent heme degradation into bilirubin and biliverdin. B) Representative crystal violet staining images 14 days post FAC (at indicated concentrations), SA (250μM) with or without HMOX1 inhibitor SnPPIX, or products of HMOX1 dependent heme degradation Biliverdin and Bilirubin. C) Relative growth of HCT116 and SW480 cells 72h post FAC (0.5, 5mM), SA (500μM) with or without SnPPIX, Biliverdin and Bilirubin. D) Schematic showing heme dependent BACH1 mediated antioxidant response. E) qPCR showing mRNA levels of various BACH1 targets - GPX4, Slc7a11, GCLC, GCLM, NQO1, NRF2, HMOX1 and FTH in HCT116 and SW480 cells 72h FAC (250μM), SA (500μM), and heme (10μM) treatment. F) Schematic showing heme dependent ATF4 stabilization and induction of integrated stress response (ISR). G) qPCR showing mRNA levels of various heme regulated inhibitor (HRI) targets CHAC1, CHOP, ATF3, ATF4 in HCT116 and SW480 cells 72h FAC (250), SA (500μM), and heme (10μM) treatment. All data are mean ± SEM. One-way ANOVA with Tukey’s multiple comparisons test for (C), (E) and (G). ∗p < 0.05, ∗∗p < 0.01, ∗∗∗p < 0.001.

**Figure S6.**
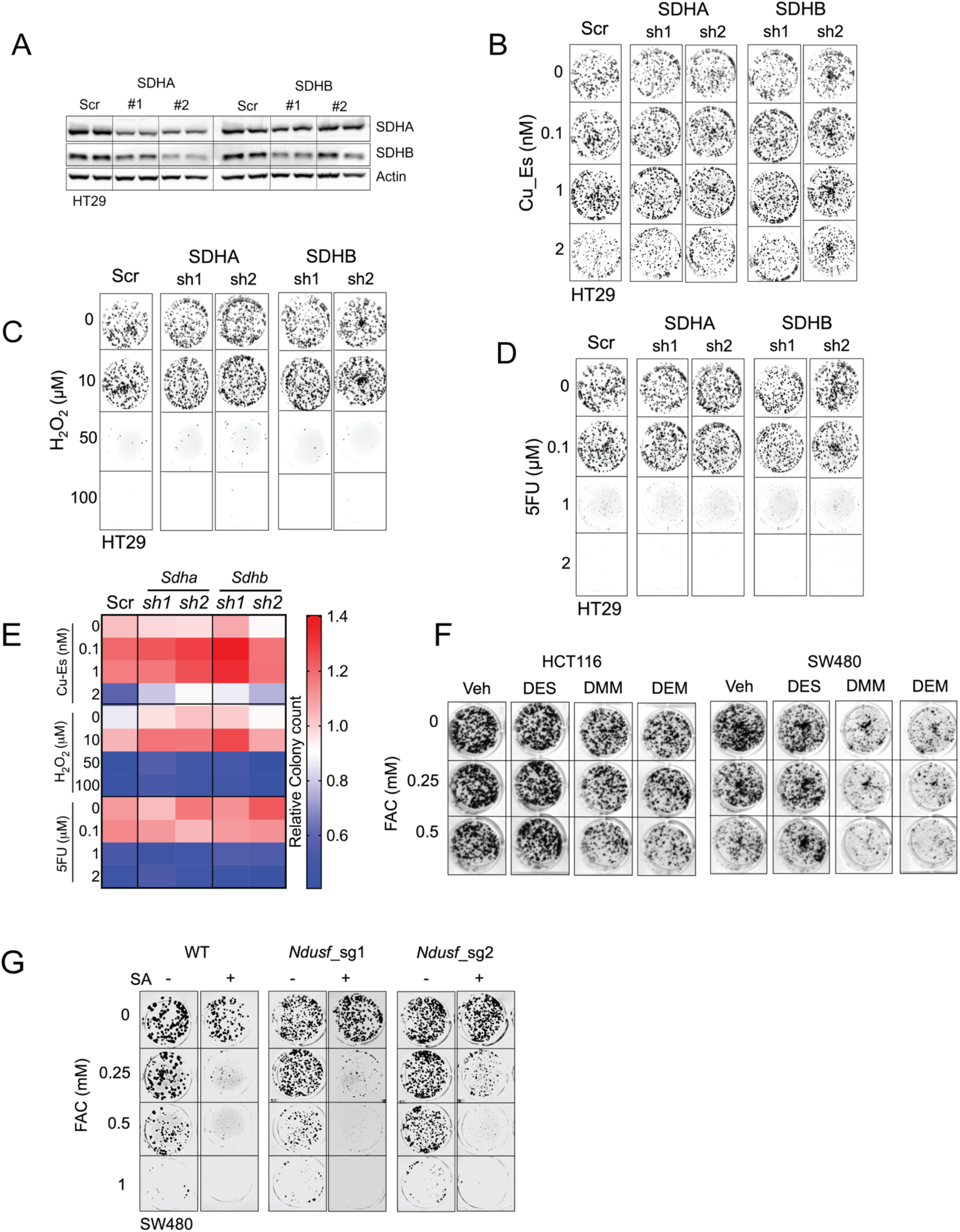
A) Western blot images showing protein expression of SDHA, SDHB and beta actin in SDHA and SDHB knockdown HT29 cells via constitutive shRNA expression. B) Representative crystal violet staining of HT29 SDHA and SDHC knockdown cells from A) 14 days post copper sulphate (100μM) + elesclomol (Cu-Es), C) hydrogen peroxide (H_2_O_2_) and D) 5-fluoroUracil (5FU) at indicated concentrations. E) Heatmap representation of relative colony count from B). F) Representative crystal violet staining images 14 days post FAC (at indicated concentrations), dimethyl succinate, dimethyl malonate, and diethyl malonate. G) Representative crystal violet staining image of NDUSF1 and 2 knockdown cells 14 days post FAC treatment (0.25, 0.5mM) with or without SA (250μM) or heme (10μM).

**Figure S7.**
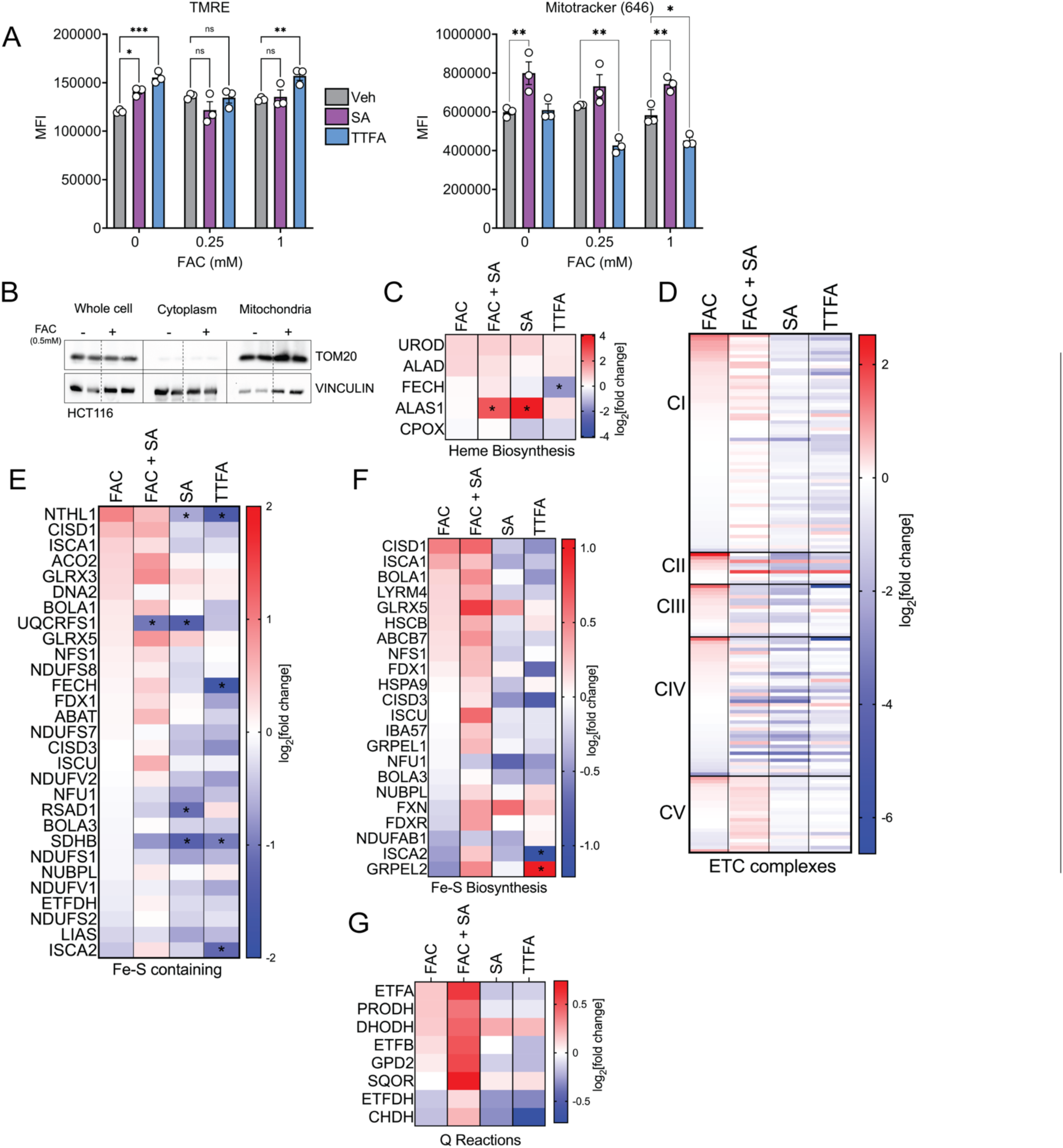
A) Mean fluorescence intensity (MFI) of Tetramethylrhodamine, ethyl ester (TRME) and mitobrilliant 646 in HCT116 cells 72h post FAC (500μM), SA (500μM) and TTFA (100μM). B) Western blot image showing protein expression of TOM20 and vinculin in the whole cell, cytoplasm and mitochondrial fractions of HCT116 cells treated with 500μM FAC for 72h. C) Heat map representation of Log2 fold change abundances for proteins involved in heme biosynthesis, D) Mitochondrial electron transport chain, E) Iron-sulfur cluster containing proteins, F) Iron-sulfur cluster biogenesis proteins, G) and Co enzyme Q reducing enzymes. * Indicates p<0.05 as compared to Vehicle treated control group. All data are mean ± SEM. One-way ANOVA with Tukey’s multiple comparisons test for (A), and t test for (D), (E), (F) and (G). ∗p < 0.05, ∗∗p < 0.01, ∗∗∗p < 0.001.

**Figure S8.**
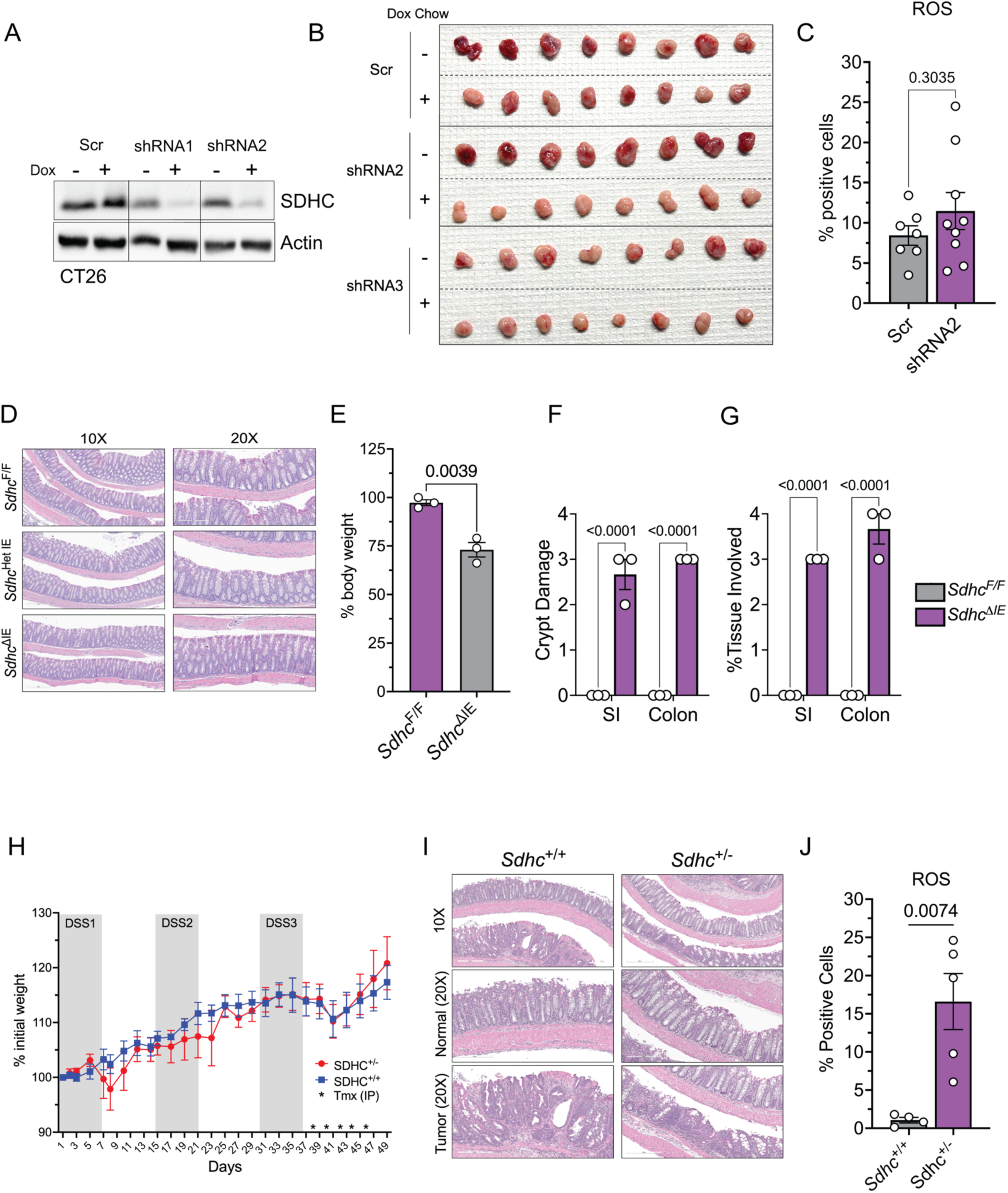
A) Western blot image showing SDHC expression and beta actin in CT26 cells stable for empty tet-pLKO-puro (Scr) or shRNA targeting SDHC (shRNA 1 or shRNA 2) 48h post Doxycycline treatment (250ng/mL). B) Gross tumor images from the xenograft tumor model with subcutaneous implantation. C) General ROS measured by free radical sensor carboxy-H2DCFDA in EPCAM positive cells from the tumors from xenograft tumor model. D) Representative H&E of small colon 5 days post tamoxifen induction via 100mg/kg body weight I.P. injection in WT (*Sdhc*^F/F^), Het (*Sdhc*^Het^ ^IE^) and KO (*Sdhc*^ΔIE^) mice. E) % body weight change in *Sdhc*^F/F^ and *Sdhc*^ΔIE^ mice fed with 350 ppm or 1000ppm iron containing chow for 7 days. F) Crypt damage score (0-4) in small intestine (SI) and colon from *Sdhc*^F/F^ and *Sdhc*^ΔIE^ mice fed with 350 ppm or 1000ppm iron containing chow for 7 days. G) % tissue involved score (0-4) in small intestine (SI) and colon from *Sdhc*^F/F^ and *Sdhc*^ΔIE^ mice fed with 350 ppm or 1000ppm iron containing chow for 7 days. H) % body weight change in *Sdhc*^F/F^ and *Sdhc*^Het^ ^IE^ mice post AOM treatment I) Representative H&E images of colon from *Sdhc*^F/F^ and *Sdhc*^Het^ ^IE^ mice on the AOM-DSS induced CAC model. J) General ROS measured by free radical sensor carboxy-H2DCFDA in EPCAM positive cells from the colonic epithelium from *Sdhc*^F/F^ and *Sdhc*^Het^ ^IE^ mice from AOM-DSS induced CAC model. All data are mean ± SEM. t test for (C), (E), (F), (G) and (J). ∗p < 0.05, ∗∗p < 0.01, ∗∗∗p < 0.001.

